# Direct and indirect neurogenesis generate a mosaic of distinct glutamatergic projection neuron types in cerebral cortex

**DOI:** 10.1101/2022.03.13.484161

**Authors:** Dhananjay Huilgol, Jesse M Levine, William Galbavy, Bor-Shuen Wang, Miao He, Shreyas M Suryanarayana, Z. Josh Huang

## Abstract

Variations in size and complexity of the cerebral cortex result from differences in neuron number and composition, which are rooted in evolutionary changes in direct and indirect neurogenesis (dNG and iNG) mediated by radial glial progenitors and intermediate progenitors, respectively. How dNG and iNG differentially contribute to cortical neuronal number, diversity, and connectivity are unknown. Establishing a genetic fate-mapping method to differentially visualize dNG and iNG in mice, we found that while both dNG and iNG contribute to all cortical structures, iNG contributes the largest relative proportions to the hippocampus and neocortex compared to insular and piriform cortex, claustrum, and the pallial amygdala. Within the neocortex, whereas dNG generates all major glutamatergic projection neuron (PN) classes, iNG differentially amplifies and diversifies PNs within each class; the two neurogenic pathways generate distinct PN types and assemble fine mosaics of lineage-based cortical subnetworks. Our results establish a ground-level lineage framework for understanding cortical development and evolution by linking foundational progenitor types and neurogenic pathways to PN types.

**Highlights:**

- A genetic strategy for differential visualization of direct and indirect neurogenesis in the same animal.
- dNG and iNG differentially contribute to piriform cortex, basolateral amygdala, hippocampus, and neocortex
- Whereas dNG generates all major PN classes, iNG differentially amplifies and diversifies PNs within each class
- dNG and iNG construct distinct cortical projection subnetworks.

## Introduction

The cerebral cortex is the largest brain structure in mammals comprising vast and diverse nerve cells that enable high-level brain functions, but the developmental mechanisms and logic underlying its neuronal diversity remain poorly understood. Cortical development begins with neurogenesis from progenitors lining the embryonic cerebral ventricle wall, which undergoes two fundamental forms of cell division that give rise to all glutamatergic neurons (Cardenas and Borrell, 2020). In direct neurogenesis (dNG), a radial glial cell (RG) undergoes asymmetric division to self-renew as well as generate one neuronal progeny (Miyata et al., 2001; Noctor et al., 2001; Rakic, 2009; Tamamaki et al., 2001); in indirect neurogenesis (iNG), RG asymmetric division produces an intermediate progenitor (IP), which then undergoes symmetric division to generate two neurons (Haubensak et al., 2004; Kriegstein et al., 2006; Miyata et al., 2004; Noctor et al., 2004). Whereas dNG is ubiquitous along the neural tube that gives rise to the central nervous system, iNG is restricted to the telencephalon giving rise to the forebrain, especially the cerebral cortex (Haubensak et al., 2004). Across evolution, while RG-mediated dNG originated before the dawn of vertebrates and has been conserved ever since, IP-mediated iNG is thought to have emerged in the last common ancestors (LCA) of amniotes and subsequently diverged along two different evolutionary paths (Cardenas and Borrell, 2020). Along the Sauropsids clade, dNG has dominated neuronal production across different pallial structures, including the 3-layered dorsal cortex of extant non-avian reptiles and the pallia of most avian species; iNG has remained rudimentary in most Sauropsids, only to expand in certain birds (corvids) where it drives increased neuron numbers and density in nuclear structures of their pallium (Cardenas et al., 2018) (Nomura et al., 2016; Striedter and Charvet, 2009). On the other hand, along the Synapsids path, iNG has expanded tremendously, particularly in the dorsal pallium, and is thought to drive the evolutionary innovation of a six-layered neocortex (Cheung et al., 2010; Florio and Huttner, 2014; Martinez-Cerdeno et al., 2006) (Villalba et al., 2021). While the amplification of cortical neuron production through IPs is inherent to iNG (Kriegstein et al., 2006), how dNG and iNG coordinate to generate the increasing diversity of glutamatergic PN types that assemble cortical networks has remained unknown.

Across the embryonic pallial subdivisions, the medial domain gives rise to the hippocampal formation; dorsal domain to the neocortex; lateral domain to insular cortex and claustrum; and the ventral domain to the piriform cortex and the pallial amygdala (Cardenas and Borrell, 2020). Among these, the six-layered neocortex comprises hierarchically organized pyramidal neuron (PyN) classes, each containing multiple finer-grained molecular and projection defined subtypes (Harris and Shepherd, 2015; Tasic et al., 2018). Within this hierarchy, the intratelencephalic (IT) class mediates myriad processing streams within the cerebral hemisphere (including ipsi- and contra-lateral intracortical and striatal projections), and the extratelencephalic (ET) class mediates subcortical outputs, including pyramidal tract (PT) neurons that project to all subcortical targets and the corticothalamic (CT) neurons that exclusively target the thalamus (Harris and Shepherd, 2015). A major unresolved question is how dNG and iNG contribute to the generation of different genetic and projection defined PyN types – the basic elements of neocortical circuit assembly and function. Furthermore, a quantitative assessment of dNG and iNG contribution to the broadly defined pallial/cortical structures and associated cytoarchitectures have not been achieved. Addressing these questions requires a method to distinguish dNG and iNG and track their developmental trajectories from progenitor types to PN types in the same animal.

Here, we deploy a novel genetic fate-mapping method to simultaneously visualize dNG and iNG as well as their PN progeny in mature cortex in mice. We have previously systematically generated mouse genetic tools targeting RG, IP, and PN types (Matho et al., 2021). Here we establish a genetic intersection-subtraction strategy and demonstrate that while dNG and iNG generate PNs for all cortical structures, iNG makes increasing contributions to cortical structures along the ventral-dorsal-medial axis, with the largest contributions to the neocortex and hippocampus. Within the neocortex, while dNG generates all major IT, PT, and CT classes, iNG differentially amplifies and diversifies PyN types within each class, with disproportionally large contribution to the IT class. Importantly, dNG and iNG derived PyN subtypes across as well as within genetically defined major subpopulations show distinct projection patterns, suggesting that they assemble fine mosaics of lineage-specified and evolutionarily-rooted cortical subnetworks. Our results reveal a ground level lineage basis of cortical development and evolution by linking foundational progenitor types and neurogenic mechanisms to PN types and their connectivity.

## Results

### A genetic strategy for differential labeling of dNG and iNG in the same animal

To distinguish and differentially fate map dNG and iNG in the same animal, we designed a genetic intersection and subtraction (IS) strategy in mice (Fig 1A). As all IPs are defined by expression of the T-box transcription factor *Tbr2* (Hevner, 2019), we generated a *Tbr2-2A-Flp* gene knockin driver line, orthogonal to multiple Cre driver lines that target RGs and PNs (Franco et al., 2012; Matho et al., 2021). Similar to our *Tbr2-2A-CreER* driver (Matho et al., 2021), the *Tbr2-2A-Flp* driver recapitulates the endogenous expression of TBR2 as early as E10.5 (Fig S1A-F) and specifically marked IPs and their PN progeny across all cortical structures (Fig S1G-J). Within the neocortex, *Tbr2-2A-Flp* marked PNs across layers, including the IT, PT, and CT classes (Fig S1H-J). Thus the *Tbr2-2A-Flp* driver enables combinatorial fate mapping of iNG and dNG with appropriate Cre driver lines and a *Intersection/Subtraction (IS)* reporter line (Fig.S2A-C); whereas Cre expression in RGs allows tracking the developmental trajectories of dNG- and iNG-derived PNs (Fig.S2B), Cre expression in postmitotic PNs can resolve their dNG or iNG origin (Fig.S2C).

**Figure 1:**
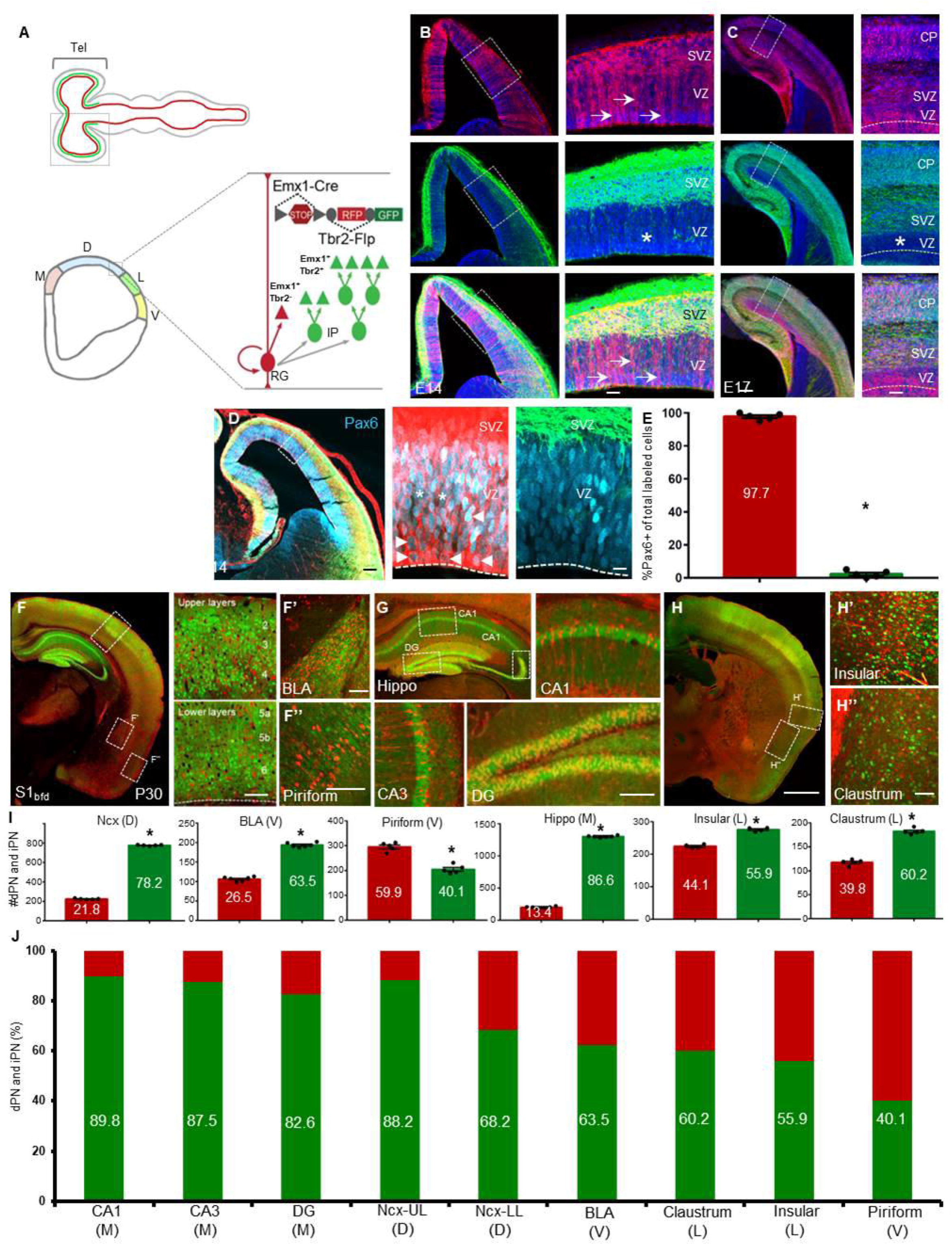
dNG and iNG differentially contribute to PNs across cortical structures (A) Top schematic shows that whereas RGs (red) exist throughout the neural tube, IPs (green) are present only in the telencephalon. Bottom coronal view from the boxed region of telencephalon shows the four subdivisions of the pallial neuroepithelium along the medio-lateral axis; medial (M), dorsal (D), lateral (L) and ventral (V) pallia, each generating distinct cortical structures. Within the neuroepithelium, RGs mediate direct neurogenesis (red) and through IPs (green), indirect neurogenesis to produce PNs (triangles). dNG and iNG can be simultaneously visualized by a genetic fate-mapping scheme using the IS reporter with *Emx1-Cre* (RG) and *Tbr2-Flp* (IP) drivers: dNG (*Emx1^+^*/*Tbr2^-^*) is labeled by RFP through ‘Cre-NOT-Flp’ subtraction; iNG (*Emx1^+^*/*Tbr2^+^*) is labeled by EGFP through ‘Cre-AND-Flp’ intersection. (B) Coronal hemi-sections of the pallial neuroepithelium, showing the labeling of RGs (top) and IPs (middle) and merged image (bottom) at E14. Right panels are magnified views of boxed regions in left panels. Arrows indicate RGs soma and radial fibers in the VZ; IPs reside in the SVZ and are absent in the VZ (asterisk). (C) Coronal hemi-sections at similar levels as in (B), but at E17. Note the appearance of dNG (RFP) and iNG (GFP) derived PNs in the cortical plate (CP). (D) Coronal hemi-sections similar to (B) showing immunohistochemistry with anti-Pax6 antibodies (cyan). Right panels are magnified views of the inset showing labeling of Pax6 with RGs (left) and IPs (right). Arrowheads indicate colocalization of Pax6 and RFP. Asterisks indicate Pax6(+) cells that are not RFP(+). (E) Quantification, 97.7% cells of the total number of labeled cells that colabel with anti-Pax6 antibody are RFP(+). (F) Coronal section of the cortex shows both dPNs (RFP) and iPNs (GFP) across laminae (upper and lower layers magnified in right panels), in the BLA (F’) and piriform cortex (F’’). (G) Coronal view of the hippocampus shows a large contribution from iPNs in different subfields; CA1, CA3 and DG. (H) Anterior coronal section shows dPNs and iPNs in the insular cortex (H’) and claustrum (H’’). (I) Quantification of differential contributions of dNG and iNG across distinct pallial structures; Y-axes are numbers of PNs quantified. Percentage of dPNs and iPNs indicated in the bar graph for respective structures shown in (F-H). (J) Quantification of differential contributions of dNG and iNG across distinct pallial structures reveal a gradient of iNG contribution from medial-to-ventral structures; percentage of iPNs are indicated in the bar graph. Note the high contribution of iPNs to the hippocampus and neocortex and their decrease in cortical structures along the medio-ventral axis. 300-1000 cells were counted in 4-6 mice for each structure. In (B-D), the dashed line indicates the ventricle boundary. Mean values are number of cells ± SEM. For (E), *P<0.0001 (compared to RFP cells), unpaired Student’s t-test. For (J), Ncx, BLA, Pir, Hippo, Ins and Cl, *P<0.0001 (compared to dPN), unpaired Student’s t-test. For (K), all structures, *P<0.0001 (compared to dPN), unpaired Student’s t-test. Scale bars 100μm (B-D); 20μm (insets B-D); 1mm (G,H); 100μm (all other scale bars). Abbreviations: tel, telencephalon; RG, radial glial cell; IP, intermediate progenitor; VZ, ventricular zone; SVZ, subventricular zone; S1_bfd_, primary somatosensory barrel field cortex; BLA, basolateral amygdala; Hippo, hippocampus; DG, dentate gyrus; Ncx, neocortex.

We first combined *Tbr2-2A-Flp* with a *Emx1-Cre* driver (Gorski et al., 2002) that targeted RGs and a *IS* reporter, which expressed RFP in Cre-NOT-Flp cells and GFP in Cre-AND-Flp cells (Fig S2A-C; He et al., 2016; Matho et al., 2021). This strategy enabled differential labeling of dNG and iNG and their derived PN progeny in the same mouse. At E14, RFP-labeled RGs resided in the VZ, characterized by their end-feet at the ventricle wall and radial fibers extending to pial surface (Fig 1B). In contrast, GFP-labeled cells were almost exclusively restricted to the SVZ with only very sparse labeling in the VZ (Fig 1B,C; Fig.S1B-E). By E17, in addition to RFP-labeled RGs and GFP-labeled IPs, dNG- and iNG-derived PNs were differentially labeled in the cortical plate (CP; Fig 1C). RG markers, PAX6 (Fig 1D-F), SOX2 and NESTIN (Fig S3) showed specific colocalization with majority of RFP-labeled cells indicating their RG identity.

### dNG and iNG differentially contribute to distinct pallial structures

To fate map dNG- and iNG-derived PNs, we first quantified the percentage of RFP and GFP labeled neurons across multiple cortical regions in P30 mice. This analysis provides the first quantitative assessment of dNG and iNG contributions across cortical structures. Consistent with previous results (Kowalczyk et al., 2009), dNG- and iNG-derived PNs constituted 21.8% and 78.2%, respectively, in all neurons of the neocortex (Fig.1I). iNG contributed more to upper layer PNs (Layers 2-4) with 11.8% dPNs and 88.2% iPNs, compared to 31.8% dPNs and 68.2% iPNs in lower layers (Layers 5-6; Fig.1J). To substantiate this result, we further used the same IS strategy with the *Lhx2-CreER* or *Fezf2-CreER* drive lines by tamoxifen induction at E12.5, which may label a different set of RGs (Matho et al., 2021). We found that 86.7% and 82.8% PNs are produced from iNG in the *Lhx2-CreER* or *Fezf2-CreER* drivers, respectively (Fig S2). Beyond the neocortex, iNG contributed to a significantly and progressively smaller fraction to the basolateral amygdala (BLA), claustrum, insular and piriform cortex (Fig.1F,H-J), consistent with the higher proportion of cycling IPs in the dorsal pallium compared to the lateral and ventral pallia (Moreau et al., 2021). Surprisingly, iNG makes the largest contribution to PNs in the hippocampus, with significantly larger fractions than in neocortex (89.9%, 87.55, 82.6% in CA1, CA3, and dentate gyrus (DG), respectively) (Fig.1G,I,J). Therefore, while both dNG and iNG contribute to all cortical structures, iNG makes larger contribution to more recently evolved structures, with disproportionate contribution to the neocortex and hippocampus. Notably, iNG contributes to cerebral structures of diverse cytoarchitectures, from six-layered neocortex, folded sheet of hippocampus, and nuclear structure of amygdala and claustrum. The fact that the hippocampus contains the largest fraction of iNG-derived PNs suggests that increased iNG per se might not have directly led to the six-layered cytoarchitecture seen in the neocortex.

### iNG amplifies and diversifies neocortical PyN types

Within the neocortex, dNG and iNG both generated all major projection classes, including IT, PT, CT (Fig S3). We thus assessed the contribution of iNG to the generation of these major PN classes. Using the *Tbr2-2A-Flp* mice in which all iNG-derived PNs expressed RFP, we quantified the percentage of RFP cells in a set of lineage transcription factor (TF)-defined PN subpopulations by immunofluorescence (Fig 2A). As expected, the vast majority of SATB2 and CUX1 IT neurons, especially those in upper layers, were derived from iNG (Fig 2B). Interestingly, half of the CTIP2 defined PT neurons derived from iNG (Fig 2C). Notably, the large majority (∼70%) of CT neurons defined by TBR1 derived from dNG; and within CT neurons, nearly 80% of the FOXP2 subpopulation and the entire TLE4 subpopulation derived from dNG (Fig 2D,E). Therefore, although dNG is initiated from the beginning of neurogenesis and generates predominantly deep layer CT and PT neurons, it continues to generate some upper layer IT neurons during late neurogenesis. Similarly, although iNG is known to generate the vast majority of upper layer IT neurons during mid-to-late neurogenesis (Mihalas et al., 2016), it also makes significant contributions to the early generation of L6 CT and L5 PT neurons.

**Figure 2:**
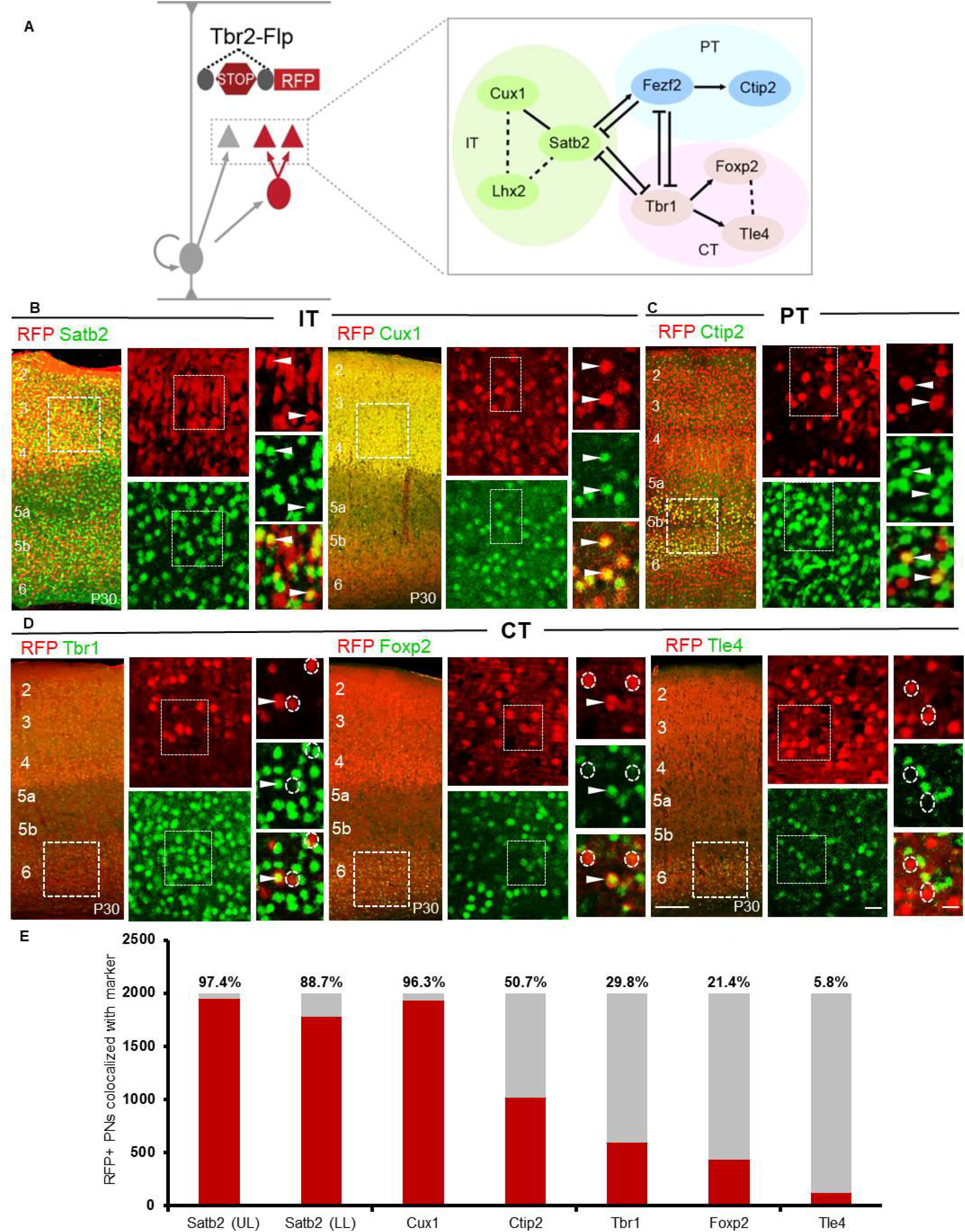
iNG differentially contributes to marker-defined cortical PN classes (A) Schematic showing a genetic strategy to label all IPs and their derived iPNs using *Tbr2-Flp* and a *Flp*-dependent reporter (also see Fig S1A). Transcriptional network interactions implicated in the postmitotic specification of IT, PT and CT PNs are also shown. (B) Representative images of immunohistochemistry using antibodies against the TFs in *Tbr2-Flp* brains: anti-SATB2 (left) and anti-CUX1 (right) label IT PNs; (C) anti-CTIP2 labels PT PNs; and (D) anti-TBR1 (left), anti-FOXP2 (middle), and anti-TLE4 (right) label CT PNs. High magnification views are from insets in low magnification images (left) for each marker. Arrowheads indicate double-positive cells; Dashed circles show non-colocalized RFP^+^ cells. (E) Quantification of immunohistochemical markers that label TF-defined iPN types in P30 *Tbr2-Flp* mice. Percentages of iPNs positive for a given TF marker are indicated above each bar graph. Quantifications were performed in S1_bfd_ from 6 sections (2000 cells) from 4-6 mice each.LMean values are number of marker-positive cells ± SEM. For Satb2, Cux1, Tbr1, Foxp2, Tle4, *P<0.0001 (compared to marker-negative RFP cells) unpaired Student’s t-test. For Ctip2, *P(n.s.) = 0.537 (compared to Ctip2-negative RFP cells) unpaired Student’s t-test. Scale bars 100μm for low magnification images (left), 20μm for high magnification views (right). Abbreviations: IT, intratelencephalic; PT, pyramidal tract; CT, corticothalamic; UL upper layer; LL, lower layer; S1_bfd_, primary somatosensory barrel field cortex.

To substantiate the above result, we deployed our genetic intersection-subtraction strategy as it can be used to resolve the dNG or iNG origin of mature PN populations (Fig S2C). By combining *Tbr2-Flp* and *IS* with a set of gene knock-in Cre driver lines that define PN subpopulations (Matho et al., 2021) (Fig 3A), we simultaneously visualized the distribution and morphology of dNG- and iNG-derived PNs within each subpopulation in the same animal (Fig 3B). Within the IT class, we have previously shown that *Cux1* positive PNs (PNs^Cux1^) mainly project within the cortex but not to the striatum, while PNs^PlxnD1^ project to ipsi- and contra-lateral cortex and striatum (Matho et al., 2021). IS labeling by postnatal tamoxifen induction in *Cux1-CreER* and *PlxnD1-CreER* drivers revealed that all postnatal PNs^Cux1^ and PNs^PlxnD1^ were GFP^+^ and thus derived from iNG (Fig 3C,D). Interestingly, early postnatal expression of *Lhx2* defines a subset of upper layer IT PNs (Matho et al., 2021), and IS labeling by P3 induction in the *Lhx2-CreER* driver revealed that 23.5% of PNs^Lhx2^ derived from dNG and 76.5% from iNG. dPNs^Lhx2^ and iPNs^Lhx2^ were extensively intermixed across L2/3 (Fig 3E).

**Figure 3:**
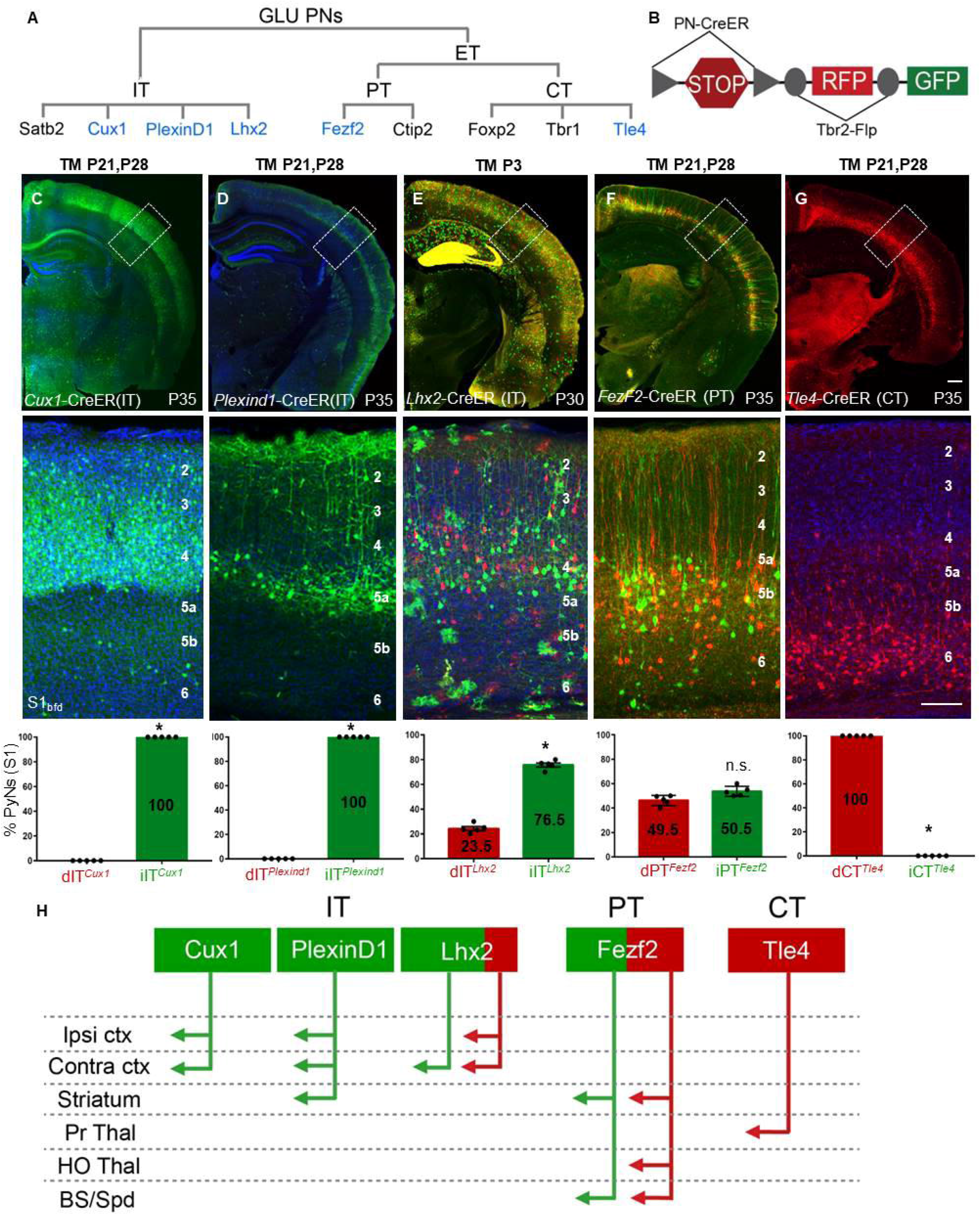
dNG and iNG differentially contribute to neocortical PN projection types (A) GLU PyNs are subdivided into broad IT and ET classes, and ET consists of PT and CT subclasses. Each of these major classes comprises multiple subpopulations defined by marker gene expression. Genes used for generating CreER driver lines are in blue. (B) Different *PN-CreER* driver lines, when combined with IS reporter line *Tbr2-Flp,* can simultaneously resolve dNG (Cre-NOT-Flp) and iNG (Cre-AND-Flp) derived subpopulations within a marker gene defined PN type. (C) PNs*^Cux1^* (L2-4 ITs, cortico-cortical PNs) and (D) PNs*^PlexinD1^* (subset of L2-5a ITs, cortico-cortical, corticostriatal PNs) were entirely iNG-derived, when corresponding driver lines were induced at P21. (E) PNs*^Lhx2^* (L2-4 ITs) were predominantly generated from iNG (76.5%) when the corresponding driver line was induced at P3. (F) PNs*^Fezf2^* (PTs) were generated equally from dNG and iNG when the driver line was induced at P21. (G) PNs*^Tle4^*, a CT subpopulation, were born entirely from dNG. (C-G) Quantifications performed for differential distribution across dNG and iNG (bottom row). Percentage of dPNs and iPNs indicated on the bar graphs. (H) dNG (red) and iNG (green) generate distinct genetic and projection defined PN subpopulations across IT, PT, and CT classes. Quantifications (C-G, bottom row) were performed in S1_bfd_ from 5 mice for 750-1500 cells each. Data are mean ± SEM. *P<0.001 (all) except, P(n.s.) = 0.0204 (PNs*^Fezf2^*), unpaired Student’s t-test. Scale bars, 1mm (low mag); 100μm (high mag). Abbreviations: GLU, glutamatergic pyramidal neurons; IT, intratencephalic, ET, extratelencephalic, PT, pyramidal tract; CT, corticothalamic; S1_bfd_, primary somatosensory barrel field cortex; Ipsi, ipsilateral; Contra, contralateral; ctx, cortex; Pr Thal, primary thalamus; HO Thal, higher order thalamus; BS, brainstem; Spd, spinal cord; TM, tamoxifen induction

Within the PT class, FEZF2 is a master TF that specifies the postmitotic PT fate, and our *Fezf2-CreER* driver captures the large majority of PT PNs (Matho et al., 2021). IS labeling by postnatal induction in *Fezf2-CreER* revealed that PNs^Fezf2^ were equally generated from dNG and iNG, and dPNs^Fezf2^ and iPNs^Fezf2^ were extensively intermixed across L5B and L6 (Fig 3F). Finally, the *Tle4-CreER* driver captures a subset of CT PNs, and IS labeling by postnatal induction in *Tle4-CreER* revealed that all PNs^Tle4^ were generated from dNG (Fig 3G). Together, these results demonstrate that dNG and iNG generate distinct subpopulations of PNs within each major class, which project to distinct cortical and subcortical regions (Fig 3H). Compared with primary somatosensory barrel field (S1_bfd_) (Fig 3), similar proportion and distribution of dPN and iPN was observed across different cortical areas including the medial prefrontal cortex (mPFC), primary motor area (M1), and primary visual area (V1) using the constitutive *Emx1-Cre* driver and the *Lhx2-CreER* (IT) and *Fezf2-CreER* (PT) drivers with postnatal TM induction (Fig S4). Therefore, iNG disproportionally diversifies IT over PT and CT subcategories and differentially diversifies genetically defined subpopulations particularly within the IT subclass across the neocortex (Figs 2E, 3H).

Beyond the neocortex, IS labeling also revealed dPNs^Fezf2^ and iPNs^Fezf2^ in the BLA, subiculum, and DG in the hippocampus. In addition, dPNs^Lhx2^ and iPNs^Lhx2^ were labeled in the DG in roughly equal ratios revealing the contribution of iNG to postnatal DG development, as previously shown (Hodge et al., 2013). An equal contribution from dNG and iNG suggests the importance of both neurogenic pathways in creating a mosaic of dentate granule cells (Fig S6). Together, these results suggest a role of both dNG and iNG in the development of PNs^Fezf2^ and PNs^Lhx2^ subpopulations in other cortical structures.

### dNG and iNG assemble distinct projection subnetworks

The extensive intermixing of dPN^Fezf2^ with iPN^Fezf2^ and dPN^Lhx2^ with iPN^Lhx2^ further raises the question of whether these lineage-distinct subpopulations represent separate subtypes even though they appear similar in laminar position and dendritic morphology. We thus examined whether these subpopulations show differences in their projection patterns. Across their subcortical targets, dPN^Fezf2^ and iPN^Fezf2^ axons remained extensively intermixed, with no clear evidence of targeting distinct regions (Fig S5B). To examine whether dPNs^Fezf2^ and iPNs^Fezf2^ differentially project to specific subcortical targets, we injected a retrograde tracer CTB into several of their targets in postnatal induced *Fezf2-CreER;Tbr2-Flp;IS* mice (Figs 4A, S7A,B). dPNs^Fezf2^ and iPNs^Fezf2^ in S1_bfd_ projected largely equally to the spinal cord (47.9% and 52.1%, respectively) and striatum (49.1% and 50.9%, respectively) (Fig S7C-G). However, of the CTB and RFP/GFP double labeled PNs, three times more dPNs^Fezf2^ (RFP) than iPNs^Fezf2^ (GFP) in S1_bfd_ somatosensory (76.2% and 23.8%, respectively) and CFA motor cortex (75.4% and 24.6%, respectively) projected to the higher order thalamic nucleus (Posterior, Po nucleus) (Fig.4C-F). PNs^Lhx2^ projected to the corpus callosum but only sparsely to the striatum (Fig S8B, C). To examine potential projection differences between dPNs^Lhx2^ and iPNs^Lhx2^, we injected CTB in the contralateral S1_bfd_ (contraS1) or ipsilateral M2 (ipsiM2) for analysis in the ipsiS1_bfd_ of P3 induced *Lhx2-CreER;Tbr2-Flp;IS* mice (Fig 4B). contraS1 received projections from a similar proportion of dPNs*^Lhx2^* and iPNs*^Lhx2^* in homotypic ipsiS1_bfd_ (Fig.4G-I), as well as in heterotypic ipsilateral M1, M2 and V1 (Fig S8D-G). In sharp contrast, ipsiM2 received a 9.4-fold higher projection from dPNs^Lhx2^ than from iPNs^Lhx2^ in ipsiS1_bfd_ (Fig 4J-L), and this dPNs^Lhx2^ versus iPNs^Lhx2^ projection difference is 12-fold higher in ipsiM1 and 9.23-fold higher in ipsiS1_fl_ (Fig S8H-K). In summary, dPNs^Lhx2^ extend much stronger projections to ipsilateral cortical areas compared to iPNs^Lhx2^. Therefore, even within the same TF-defined subpopulations that are highly intermixed, dPNs and iPNs show preferential projection patterns (Fig. 4F,M). Together with the categorical distinction of dNG-generated PNs^Tle4^ and iNG-generated PNs^Cux1^ and PNs^PlxnD1^, these results indicate that dNG and iNG generate distinct projection subtypes within marker defined PN subpopulations.

**Figure 4:**
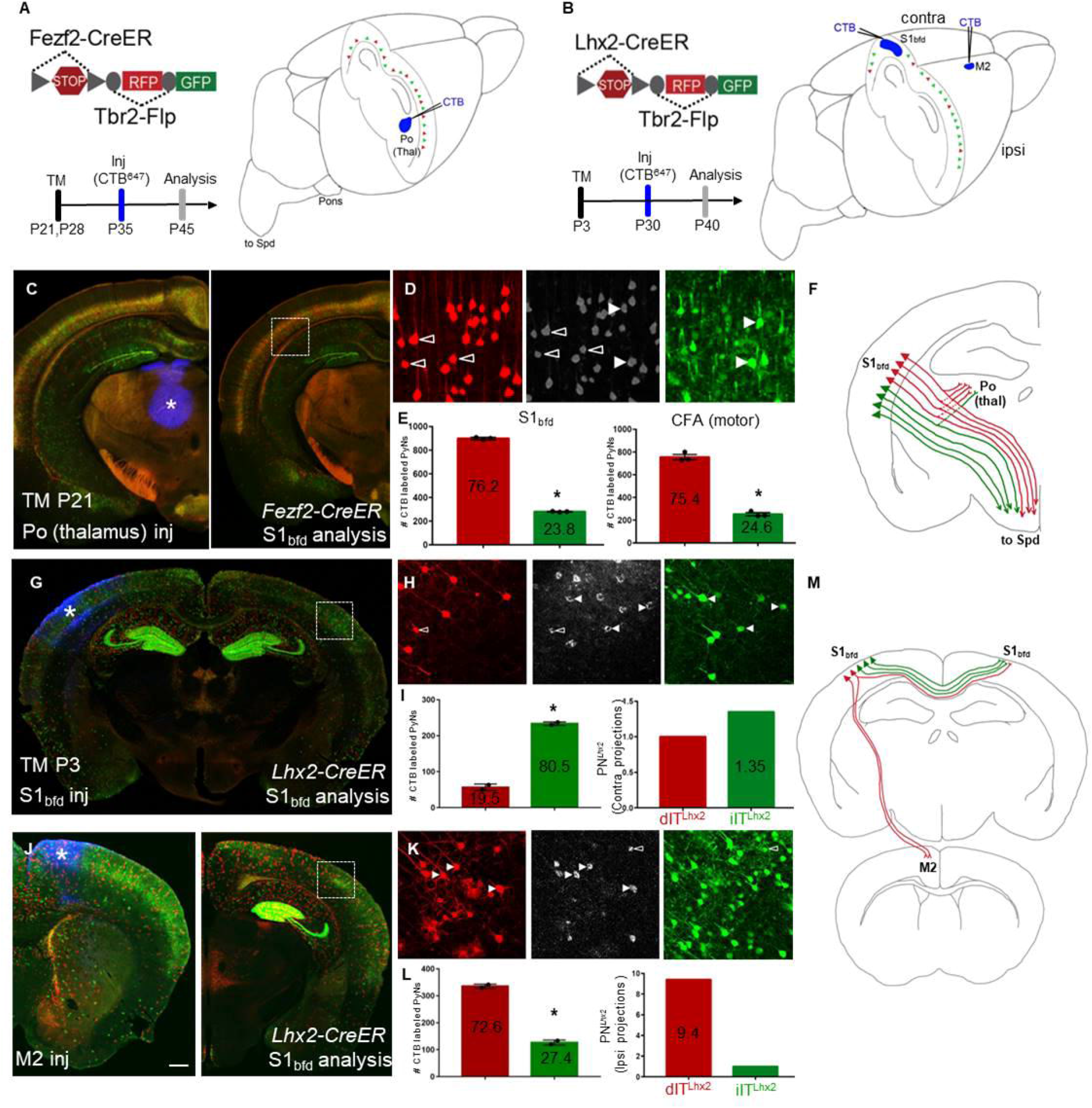
dPNs^Fezf2^ and dPNs^Lhx2^ in the neocortex project preferentially to higher-order thalamus and ipsilateral cortical areas, respectively (A-B) Schematics depicting retrograde CTB labeling from the Po (higher order) nucleus of thalamus in *Fezf2-CreER;Tbr2-Flp;IS* (PNs^Fezf2^) mice induced at P21(A) or from either S1_bfd_ or M2 in *Lhx2-CreER; Tbr2-Flp;IS* (PNs^Lhx2^) mice induced at P3 (B). (C) Coronal hemisection of the neocortex from a PNs^Fezf2^ brain showing the injection site, Po (asterisk, left) and analysis in S1_bfd_ (right). (D) CTB labeling (middle panel) colocalized with dPNs^Fezf2^ (open arrowheads, left panel) or iPNs^Fezf2^ (white arrowheads, right panel). (E) Quantification in S1_bfd_ (left) showed 76.2% of CTB and RFP/GFP double labeled cells were dPNs^Fezf2^ and 23.8% were iPNs^Fezf2^. In the CFA (motor area), 75.4% of CTB and RFP/GFP double labeled cells were dPNs^Fezf2^ and 24.6% were iPNs^Fezf2^. (F) Schematic showing that dPNs^Fezf2^ preferentially project to the higher-order thalamus when compared with iPNs^Fezf2^. (G) CTB injected in S1_bfd_ (asterisk) of PNs^Lhx2^ mice and analyzed for colocalization in the contraS1_bfd_. (H) CTB (middle panel) colocalizes with more iPNs^Lhx2^ (right panel arrowheads, GFP) and relatively fewer dPNs^Lhx2^ (left panel open arrowheads, RFP) in contraS1_bfd_. (I) Quantification shows that 80.5% of CTB-XFP double labeled cells were iPNs^Lhx2^ and 19.5% were dPNs^Lhx2^ (left). When normalized to the ratio of dPNs^Lhx2^ and iPNs^Lhx2^, iPNs^Lhx2^ showed 1.35 fold more than dPNs^Lhx2^ in projection to contrS1. (J) CTB injected in M2 (asterisk, left) and analyzed in PNs^Lhx2^ mice in the ipsilateral, ipsiS1_bfd_ (right) (K) CTB colocalizes with more dPNs^Lhx2^ (left panel white arrowheads, RFP) compared to iPNs^Lhx2^ (right panel open arrowheads, GFP). (L) Among CTB and RFP/GFP double labeled cells in ipsiS1_bfd_, 72.6% were dPNs^Lhx2^ and 27.4% were iPNs ^Lhx2^ (left). When normalized to the ratio of dPNs^Lhx2^ and iPNs^Lhx2^ in ipsiS1_bfd_, dPNs^Lhx2^ showed a 9.4-fold higher projection to iM2 than iPNs^Lhx2^. (M) Summary schematic showing dPNs^Lhx2^ preferentially projecting to ipsilateral cortical areas when compared with iPNs^Lhx2^. Quantifications were performed in S1_bfd_ or CFA from 1000 cells, 3-4 mice for PNs^Fezf2^ in (E). Mean values are number of CTB labeled PNs ± SEM *P<0.0001 unpaired Student’s t-test (compared to dPN), unpaired Student’s t-test for both S1_bfd_ and CFA. For PNs^Lhx2^, 300-450 cells were counted from S1_bfd_ in 3-4 animals in (I,L). Mean values are number of CTB labeled PNs ± SEM *P<0.0001 unpaired Student’s t-test (compared to dPN). Scale bars, low mag (C,G,J) 1mm; high mag (D,H,K) 100μm. Abbreviations: Po, posterior nucleus of thalamus; Thal, thalamus; S1_bfd_, primary somatosensory barrel field cortex; M2, secondary motor cortex; inj, injection; CFA, caudal forelimb area; Spd, spinal cord; TM, tamoxifen induction.

## Discussion

Our findings provide the first quantitative assessment of dNG and iNG contributions across cerebral cortical structures and to distinct PN types in the neocortex that assemble different subnetworks. Previous studies have emphasized the role of SVZ/iNG in the generation of upper layer PNs of the neocortex, suggesting that the rise of iNG in mammals contribute to the formation of a six-layered cytoarchitecture (Cardenas and Borrell, 2020; Cheung et al., 2010; Martinez-Cerdeno et al., 2006; Villalba et al., 2021). Our results demonstrate that iNG in fact contributes to the generation of all pallial/cortical structures in mice, including those which are considered phylogenetically “old”, archi- and paleo-cortices. We provide the first quantitative assessment of dNG and iNG contribution across these structures, from the laminated neocortex, hippocampus and piriform cortex to nuclear structures of the amygdala and claustrum. It is interesting to note that, beyond mammals, the increase of iNG in corvids correlates with the rise of laminated (Wulst/hyperpallium) and nuclear pallial structures (DVR) (Cardenas et al., 2018; Nomura et al., 2016; Striedter and Charvet, 2009). We further reveal that along the cortical medial-lateral axis, iNG makes progressively lower contributions, with sharp decreases in the amygdala and piriform cortex. Surprisingly, iNG makes the largest relative contribution to the hippocampus, significantly more than the neocortex. These results suggest that the rise of iNG per se might not have simply led to increased lamination in cytoarchitecture (i.e. six-layered neocortex). More likely, the fundamental consequence of iNG is the increase in cell number and diversity, which can assemble multiple forms of cytoarchitectures ranging from a folded cell sheet of the hippocampus to six-layered neocortex and to nuclear structures like the amygdala and claustrum. Consistent with this notion, hippocampal neurogenesis proceeds in parallel with that of the neocortex (Bond et al., 2020; Chen et al., 2017; Xu et al., 2014), and recent single cell transcriptome analysis in mouse hippocampus has revealed a cell type diversity comparable to that of the neocortex (Yao et al., 2021).

A key component in neocortical development and evolution has been the diversification of PN types (Arendt et al., 2016; Briscoe and Ragsdale, 2018; Colquitt et al., 2021; Tosches et al., 2018). Although several previous studies showed that iNG generates PNs in all neocortical layers, and particularly those in the upper layers (Mihalas et al., 2016; Mihalas and Hevner, 2018; Vasistha et al., 2015), and suggested differences in dendritic arborization and electrophysiological properties between *Tbr2-* and non-*Tbr2* derived PNs within the same cortical layer (Guillamon-Vivancos et al., 2019; Tyler et al., 2015), these studies have not resolved the relative contributions of dNG and iNG to different PN types. Here we show that dNG in fact generates all major cortical PyN classes, while iNG differentially amplifies and diversifies PyN types within each class defined by projection pattern and molecular markers beyond laminar location. iNG not only makes disproportionally large contribution to the IT class as expected, it also contributes to half of the PT class and a significant portion of the CT class. Interestingly, dNG remains the major source of CT class, likely reflecting its dominance over iNG during the early phase of neurogenesis that gives rise to L6 CT neurons. It is conceivable that the CT class may have evolved in mammals from the diversification of ancestral “PT-type” cells which can be found in several vertebrates (Dugas-Ford et al., 2012; Ebbesson and Schroeder, 1971; Ocana et al., 2015).

Furthermore, dNG and iNG derived PN types across (PNs^Cux1^, PNs^PlxnD1^, PNs^Tle4^) as well as within (PNs^Fezf2^, PNs^Lhx2^) genetically defined major subpopulations show distinct projection patterns. These results indicate that dNG and iNG assemble a fine mosaic of lineage-based and likely evolutionarily-rooted cortical subnetworks (Fig. 5). As RG-dNG and IP-iNG undergo fundamentally distinct cell division patterns, their neuronal progenies derive from different birth pattern and order (asymmetric division from RGs vs symmetric cell division from IPs), which likely confer differential chromatin landscapes that impact transcription profiles (Pinson and Huttner, 2021). Multi-omics analysis of dNG and iNG-derived PNs may reveal their epigenomic and transcriptomic distinctions that underlie their phenotypic distinctions. At the level of circuit connectivity, the categorical distinction between iNG-derived PNs^Cux1^ and PNs^PlxnD1^ versus dNG-derived PNs^Tle4^ suggest separate construction of major cortical networks and associated brain systems. Our finding of seemingly more subtle projection differences between dNG- and iNG-derived PN^Fezf2^ and PN^Lhx2^ by retrograde labeling are likely underestimates; methods that quantify synaptic connectivity may reveal further distinction between dPNs and iPNs within genetically defined subpopulations. A major further challenge is to discover whether and how the distinction of dNG and iNG-derived PNs manifest at the level of circuit function underlying behavior; such studies require methods to differentially monitor and manipulate the activity of dNG and iNG-derived PNs.

**Figure 5:**
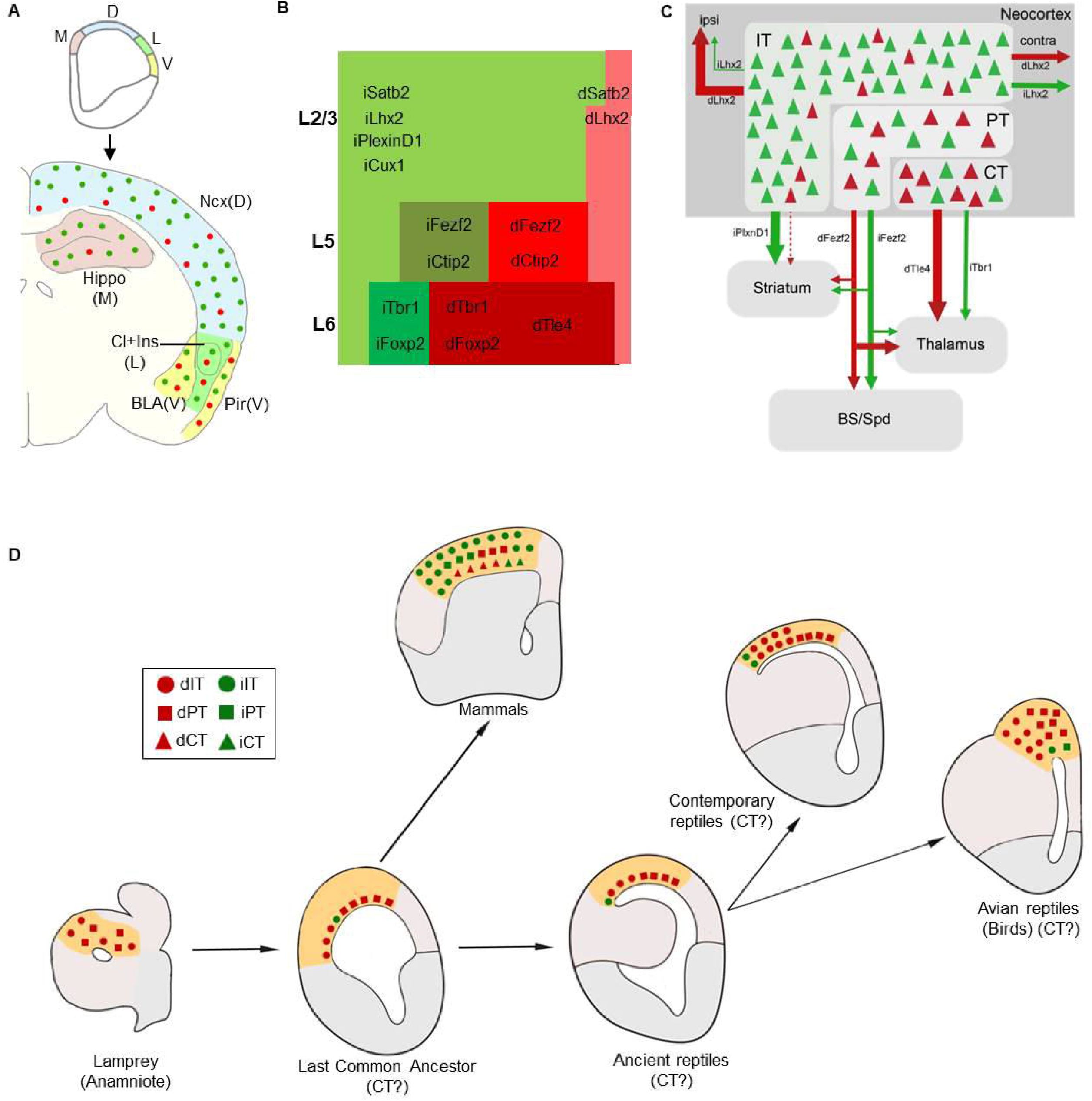
Schematics summarizing dNG and iNG contribution to cortical structures, a mosaic of neocortical PN types and subnetworks, and evolutionary implications (A) Along the medial to ventral axis of the mouse embryonic pallium, dNG and iNG generate dPNs (red) and iPNs (green) that populate all cortical structures, with decreasing iNG contributions to lateral and ventral structures. (B) Within the neocortex, dNG generates CT (dark shade), PT (medium shade), and IT (light shade) class dPNs (red) across layers, whereas iNG differentially amplifies and diversifies genetically defined iPN types (green) within each class. iPNs have a disproportionally large contribution to the IT class. (C) dNG (red) and iNG (green)-derived PN types are highly intermixed within the neocortex and yet show distinct projection patterns both across and within genetically defined subpopulations. Thus, dNG and iNG construct lineage-based fine mosaics of cortical subnetworks. (D) A conceptual schema depicting the evolutionary trajectory of dNG (red) and iNG (green) with their derived major PN types in dorsal pallial homologs across vertebrates (modified from Suryanarayana et al., 2021; Briscoe and Ragsdale, 2018). dNG and their derived IT (circle) and PT (square) classes are present in lamprey (cyclostomes) and thus predate the dawn of vertebrates. IPs and iNG may have originated in the last common ancestor of amniotes. Among the Sauropsids, dNG has dominated PN production across different pallial structures, including the three-layered dorsal cortex of extant non-avian reptiles and the pallia of most avian species; iNG has remained rudimentary, only to expand in certain birds (corvids) where it drives increased neuron numbers and density in nuclear structures of their pallium. Among Synapsids including mammals, the expansion of iNG greatly amplifies and diversifies PN types across neocortical layers and PN classes. Abbreviations: M, medial pallium; D, dorsal pallium; L, lateral pallium; V, ventral pallium; Ncx, neocortex; Hippo, hippocampus, Cl, claustrum; Ins, insular cortex; BLA, basolateral amygdala; Pir, piriform cortex; ipsi, ipsilateral; contra, contralateral, BS, brain stem; Spd, spinal cord; LCA, last common amniote, IT, intratelencephalic; PT; pyramidal tract; CT, corticothalamic.

Several intracellular and extracellular factors have been shown to influence the balance between dNG and iNG in mammals by affecting the cell division pattern of RGs (Postiglione et al., 2011; Vitali et al., 2018; Lv et al., 2019; Hasenpusch-Theil et al., 2020) and intercellularsignaling pathways (Cardenas and Borrell, 2020; Taverna et al., 2015),. In particular, the interplay of Robo and Notch signaling levels has been implicated in determining the relative proportion of dNG and iNG across amniotes (Cardenas et al., 2018). High Slit/Robo and low Dll1 signaling are necessary and sufficient to drive direct neurogenesis, suggesting that modulation in activity levels of conserved signaling pathways is likely a mechanism driving the expansion and increased complexity of the mammalian neocortex during amniote evolution (Cardenas and Borrell, 2018; 2020). Comparative studies on the molecular mechanisms of RG division patterns across amniotes will provide further insights in the evolutionary expansion of IPs and iNG.

As brain structures assemble and organize at multiple levels from molecules to cells, embryological territories, and neural circuits, these levels can evolve independently of one another, and homology at one level does not require conservation at other levels. Given that cell types are the elemental units of gene regulation as well as neural circuit assembly, they also constitute the basic units of conservation and divergence linking genomic changes to the evolutionary innovations of tissue organization and behavior. Indeed, recent studies suggest that extant amniotes possess a variety of divergent pallial structures, from six-layered neocortex in mammals to three-layered dorsal cortex in non-avian reptiles to nucleus-like pallia in birds. They share a conserved set of neuronal cell types and circuitries, the basic elements of which can be traced back even to the earliest of vertebrates (Briscoe and Ragsdale, 2018; Cardenas and Borrell, 2020; Lamanna et al., 2022; Suryanarayana et al., 2021) (Fig 5). A key approach in this cell type perspective of cortical evolution is to delineate the developmental trajectories from progenitor types to neuronal cell types in the assembly of brain circuits. Our finding of distinct developmental trajectories of dNG and iNG begin to provide a ground-level lineage framework of cortical development and evolution by linking foundational progenitor types and neurogenic pathways with conserved and diversified PN types across species, dating from the pan-vertebrate dNG to the emergence of iNG in the amniote LCA (Briscoe and Ragsdale, 2018; Cardenas and Borrell, 2020; Suryanarayana et al., 2021). Such a cell lineage framework may facilitate exploring the evolutionary origin of the neocortex and its relationship to possible homologous pallial structures across vertebrates (Suryanarayana et al., 2021). Cellular resolution multi-modal analysis based on this lineage framework may guide evolutionary comparisons, linking developmental genetic programs in progenitor types to transcriptome profiles in cell types (Colquitt et al., 2021; Tosches et al., 2018) and to neural circuit organization across cortical structures, including the neocortex.

## Acknowledgements

We thank Debra L. Silver, Richard Mooney and György Buzsáki for comments on the manuscript. We thank L. Li and Priscilla Wu at CSHL for help with generation of *Tbr2-2A-Flp* knock-in; Jonathan Werner for help with quantification related to Figs 2,4; the CSHL and Duke University animal resources for mouse husbandry; and the CSHL Microscopy shared resource and Duke University Light Microscopy Core Facility.

## Funding

NIH grant U19MH114823-01 (ZJH).

NIH Director’s Pioneer Award 1DP1MH129954-01 (ZJH).

Human Frontier Science Program long-term fellowship LT000075/2014-L (DH).

NARSAD Young Investigator Grant no. 26327 (DH).

NRSA F30 Medical Scientist Predoctoral Fellowship 5F30MH108333 (JML).

NRSA Postdoctoral Fellowship NIH5F32NS096877-03 (B-SW).

International Postdoc Grant, Swedish Research Council. Grant no: 2021-00238 (SMS).

## Author Contributions

ZJH and DH conceived the project.

ZJH, DH and JML designed the experiments,

MH designed the *Tbr2-2A-FlpO* knock-in mouse line.

DH and JML performed fate mapping, immunohistochemistry, imaging and quantification.

DH, B-SW and WG conducted injection experiments.

ZJH, DH and SMS wrote the manuscript.

## Declaration of Interests

The authors declare no competing interests.

## STAR Methods

### Resource Availability

#### Lead Contact

Further information and requests for resources and reagents should be directed to and will be fulfilled by the Lead Contact, Z. Josh Huang.

#### Materials Availability

Materials generated in this study are available on request to the Lead Contact.

#### Data and Code Availability

The datasets generated during this study are available upon request.

### Experimental Details

#### Mouse breeding

Mouse related experimental procedures were approved by the Institutional Animal Care and Use Committee (IACUC) of the Cold Spring Harbor Laboratory and Duke University in accordance with NIH guidelines. All genetically targeted animals have been backcrossed 6 generations to a Swiss-Webster background.

#### Generation of <u>Tbr2-2A-Flp</u> knock-in mouse line

*Tbr2-2A-Flp* was generated by inserting a 2A-Flp cassette in-frame before the STOP codon of the targeted gene. Targeting vectors were generated using a PCR-based cloning approach as described before (He et al., 2016; Matho et al., 2021).

#### Generation of <u>Intersection-Subtraction</u> reporter line

The *IS* reporter was generated as described previously (https://www.jax.org/strain/036760; He et al., 2016; Matho et al., 2022). Briefly, a STOP cassette flanked with *loxP* sites and dTomato (dimer) sequence flanked with *FRT* sites was targeted at the *Gt(ROSA)26Sor* locus, preventing the transcription of enhanced green fluorescent protein (EGFP). dTomato is expressed following *cre*-mediated recombination, while EGFP is expressed following both *cre*- and *flp*-mediated recombination. The targeting vector was linearized and electroporated into a 129SVj/B6 F1 hybrid ES cell line (V6.5, Open Biosystems). G418-resistant ES clones were first screened by PCR and then confirmed by Southern blotting. Positive ES cell clones were used for tetraploid complementation to obtain heterozygous male mice using standard procedures.

While recombination efficiency of all the cre-expressing lines used in this study is ∼100% (Gorski et al., 2002; Matho et al., 2021) and Tbr2-Flp provides near complete recombination, the IS reporter requires both cre and flp recombinations which may not be fully efficient. However, in this study we have assayed the relative proportion of dNG vs iNG and dPN vs iPN, not their absolute numbers. Also, tamoxifen induction of CreER recombination will inherently vary, but still report the proportion of dNG vs iNG and dPN vs iPN.

#### Tamoxifen induction

Tamoxifen (T5648, Sigma) was prepared by dissolving the powder in corn oil (20Lmg/ml) and either applying a sonication pulse for 60s or constant magnetic stirring overnight at 37 °C. A 100–200Lmg/kg dose was administered by intraperitoneal injection at the appropriate age; If two doses, 100mg/kg doses were administered at P21 and P28. For experiments with *Lhx2-CreER*, 200mg/kg was administered intraperitoneally at P3 from a diluted stock of 5mg/ml.

#### Immunohistochemistry

Adult mice were anaesthetized (using Avertin) and transcardially perfused with saline followed by 4% paraformaldehyde (PFA) in 0.1 M phosphate buffer. After post-fixation, brains were rinsed three times in PBS and sectioned at a 65-70µm thickness with a Leica VT1000S vibratome. Embryo heads were collected in PBS and fixed in 4% PFA for 4h at room temperature, rinsed three times with PBS, equilibrated in 30% sucrose-PBS, frozen in OCT compound and cut on a cryostat (Leica, CM3050S) at 25µm coronal sections. Sections were treated with a blocking solution (10% normal goat serum and 0.2% Triton-X100 in 1X PBS) for 1h, then incubated overnight at 4°C with primary antibodies diluted in the blocking solution. Sections were washed three times in PBS and incubated for 2h at room temperature with corresponding secondary antibodies, Goat or Donkey Alexa Fluor 488, 594 or 647 (1:500, Life Technologies) and DAPI to label nuclei (1:1000 in PBS, Life Technologies, 33342). Sections were washed three times with PBS and dry-mounted on slides using Fluoromount-G (SouthernBiotech, 0100-01) mounting medium.

### Primary Antibodies

Anti-GFP (1:1000, Aves, GFP-1020), anti-RFP (1:1000, Rockland Pharmaceuticals, 600-401-379), anti-SATB2 (1:20, Abcam ab51502), anti-CUX1 (1:100, SantaCruz 13024), anti-CTIP2 (1:100, Abcam 18465), anti-TBR1 (1:250, MilliporeSigma AB2261), anti-FOXP2 (1:500, Santa Cruz sc-517261), anti-TLE4 (1:300, Santa Cruz sc-365406), anti-Tbr2 (1:250, EMD Millipore AB15894), anti-Pax6 (1:300, MBL Intnl PD022), anti-Sox2 (1:300, Millipore Sigma AB5603 and anti-Nestin (1:300, Abcam ab22035) were used.

For anti-CTIP2 and anti-SATB2, brains were postfixed in 4% PFA for 4hrs at room temperature. For all other antibodies, postfixation was done overnight at 4°C

#### Imaging and Quantification

Imaging from serially mounted sections was performed on a Zeiss LSM 780 or 710 confocal microscope (CSHL St. Giles Advanced Microscopy Center and Duke University Light Microscopy Core Facility) using objectives 10x and 63x for embryos, and 5x, 10x and 20x for adult mouse brains.

All imaging was done using Zeiss LSM 710 or 780 fluorescence confocal microscopes using objectives, 5x for tilescan, 10x or 20x for z-stacks. For embryos, high magnification images were obtained using 63x oil objective. To determine colocalization in adult mouse brains, confocal z-stacks were obtained centered in S1_bfd,_ using a 20x objective. We manually determined colocalization for the desired markers by looking in individual z-planes using ImageJ/FIJI software. All quantifications were performed by two individuals. Statistics and plotting of graphs were done using GraphPad Prism 7 and Microsoft Excel 2010. For embryonic experiments (Figs 1, S1,S3), high-magnification insets are not maximum intensity projections. To observe the morphology of IPs (Fig S1) and quantification of colocalization with RG markers (Figs 1, S3), only a few sections from the z-plane in low-magnification images have been projected in the high-magnification images. For colocalization experiments with PAX6, SOX2 AND NESTIN in *Emx1-Cre; Tbr2-Flp; IS* embryonic brains, DAPI was used identify cells since the RFP labeling is across the cell body and along the radial fiber of RGs. For embryonic quantifications, we counted 70-200 cells depending on the extent of labeling from at least 5 embryonic brains across 2 litters. For all adult neocortex quantifications, we counted in 1mm x 1mm area from at least 6 sections, from 5-6 adult brains. Number of cells counted for *Emx1-Tbr2-IS* experiment: Neocortex, 1000 cells; CA1, 500 cells; CA3, 500 cells; DG, 500 cells; BLA, 300 cells; Claustrum, 300 cells; Insular cortex, 500 cells; Piriform cortex, 500 cells. For each structure we quantified at least 6 sections from 4-6 brains. To perform molecular characterization of *Tbr2-2A-Flp* brains, we stained vibratome sections for SATB2, CUX1, CTIP2, TBR1, FOXP2 and TLE4. Percentage positive cells were calculated from an average number of 2000 RFP+ cells per staining. Total number of cells counted for *PN-CreER; Tbr2-flp; IS* experiments for each line was between 750-1500. For *Fezf2-CreER; Tbr2-flp; IS* and *Lhx2-CreER; Tbr2-flp; IS* experiments, number of cells counted are: BLA, 300; Subiculum, 500; DG^Fezf2^, 150; DG^Lhx2^, 1000. For each driver line we quantified at least 6 sections from 4-6 brains. PN numbers are different due to differences in labelling density.

For CTB quantifications in Fig 4G-L and Fig S6D-K, “normalization” refers to the ratio of number of CTB/XFP double positive cells to the total number of XFP positive cells observed (XFP is is either RFP or GFP). This aided in determine the fold-difference between the projections from dPNs^Lhx2^ and iPNs^Lhx2^ relative to their total number. CTB quantifications for PNs^Fezf2^ were done from ∼1000 cells from 3-5 mice (Figs 4, S5). For PNs^Lhx2^, quantifications were done in ipsiS1_bfd_ from ∼300 cells for contraS1_bfd_ and ∼450 cells from ipsiM2 injections, from 3-4 brains each (Fig 4). In Fig S6, from contraS1_bfd_ injections, colocalization was observed in ipsiM1 (∼400 cells), ipsiM2 (∼90 cells), ipsiV1 (∼120 cells). From ipsiM2 injections, colocalization was seen in contraM1 (∼200 cells), ipsiM1 (∼300 cells) and ipsiS1_fl_ (∼500 cells).

#### Stereotaxic Injections

Adult mice were anaesthetized by 2% isofluorane inhalation with 0.41/min airflow. Preemptive analgesics, 5mg/kg ketoprofen and 0.5mg/kg dexamethasone, were administered subcutaneously before the surgery. Lidocaine (2–4Lmg/kg) was applied intra-incisionally. Mice were mounted on a stereotaxic headframe (Kopf Instruments, 940 series), and coordinates were identified. An incision was made over the scalp, a small burr hole drilled in the skull and injections were performed in either the primary somatosensory barrel field cortex (S1_bfd_):1.7 posterior relative to bregma, 3.75 lateral, 0.5-0.3 in depth or in the secondary motor cortex (M2): 1.05 anterior relative to bregma, 1.0 lateral, 0.5 in depth. A pulled glass pipette tip of 20–30Lμm containing CTB^647^ (ThermoFischer Scientific, C34778) or AAV (Addgene, AAV-PHP.eB) was lowered into the brain. A 500nl (CTB) or 300-400nl (AAV) volume was delivered at a 30nl/min using a Picospritzer (General Valve Corp); to prevent backflow, the pipette was maintained in place for 10Lmin prior to retraction. The incision was sutured with Tissueglue (3M Vetbond), following which mice were kept warm at 37°C until complete recovery.

## Supplemental Information

***Supplemental Figs S1-S8 legends***

---

**Table 1:** Resources used in the study.

| Driver lines<br>(Cre/CreER or Flp) | Reporter lines | Antibodies | Retrograde label |
| --- | --- | --- | --- |
| Tbr2-FlpO<br>(Current Study)<br>Emx1-IRES-Cre<sup>(Gorski et al., 2002)</sup><br>RRID:IMSR_JAX:005628<br>Cux1-2A-CreER<sup>(Matho et al., 2021)</sup><br>RRID:IMSR_JAX:036300<br>PlexinD1-2A-CreER<sup>(Matho et al., 2021)</sup><br>RRID:IMSR_JAX:036295<br>Lhx2-2A-CreER<sup>(Matho et al., 2021)</sup><br>RRID:IMSR_JAX:036293<br>Fezf2-2A-CreER<sup>(Matho et al., 2021)</sup><br>RRID:IMSR_JAX:036296<br>Tle4-2A-CreER<sup>(Matho et al., 2021)</sup><br>#RRID:IMSR_JAX:036298 | FSF-tdTomato<sup>(Matho et al., 2021)</sup><br>Intersection-Subtraction<br>(IS) <sup>(He et al., 2016; Matho et al., 2021)</sup><br>RRID:IMSR_JAX:028582<br>RRID:IMSR_JAX:036760 | Anti-RFP 1:1000<br>(Rockland Cat# 600-401-379,<br>RRID:AB_2209751)<br>Anti-GFP 1:1000<br>(Aves Labs Cat# GFP-1020,<br>RRID:AB_10000240)<br>Anti-Satb2 1:20<br>(Abcam Cat# ab51502,<br>RRID:AB_882455)<br>Anti-Cux1 1:100<br>(Santa Cruz Biotechnology<br>Cat# sc-13024,<br>RRID:AB_2261231)<br>Anti-Ctip2 1:150<br>(Abcam Cat# ab18465,<br>RRID:AB_2064130)<br>Anti-Foxp2 1:300<br>(Santa Cruz Biotechnology<br>Cat# sc-517261,<br>RRID:AB_2721204)<br>Anti-Tbr1 1:300<br>(Millipore Cat# AB10554,<br>RRID:AB_10806888)<br>Anti-Tle4 1:300<br>(Santa Cruz Biotechnology<br>Cat# sc-365406,<br>RRID:AB_10841582)<br>Anti-Tbr2 1:250<br>(Millipore Cat# AB15894,<br>RRID:AB_10615604)<br>Anti-Pax6 1:300<br>(MBL International Cat#<br>PD022, RRID:AB_1520876)<br>Anti-Sox2 1:300<br>(Millipore Cat# AB5603,<br>RRID:AB_2286686)<br>Anti-Nestin 1:300<br>(Abcam Cat# ab22035,<br>RRID:AB_446723) | Cholera Toxin Subunit B<br>(Recombinant), Alexa Fluor<br>647 (ThermoFischer C34778) |

## Notes

### Competing Interest Statement

The authors have declared no competing interest.

### Summary of Updates

This manuscript has been revised based on Reviewers' comments.

