## Supplemental Information for "Direct and indirect neurogenesis generate a mosaic of distinct glutamatergic projection neuron types in cerebral cortex"

**This PDF file includes:**

Supplemental Figs. S1 to S8

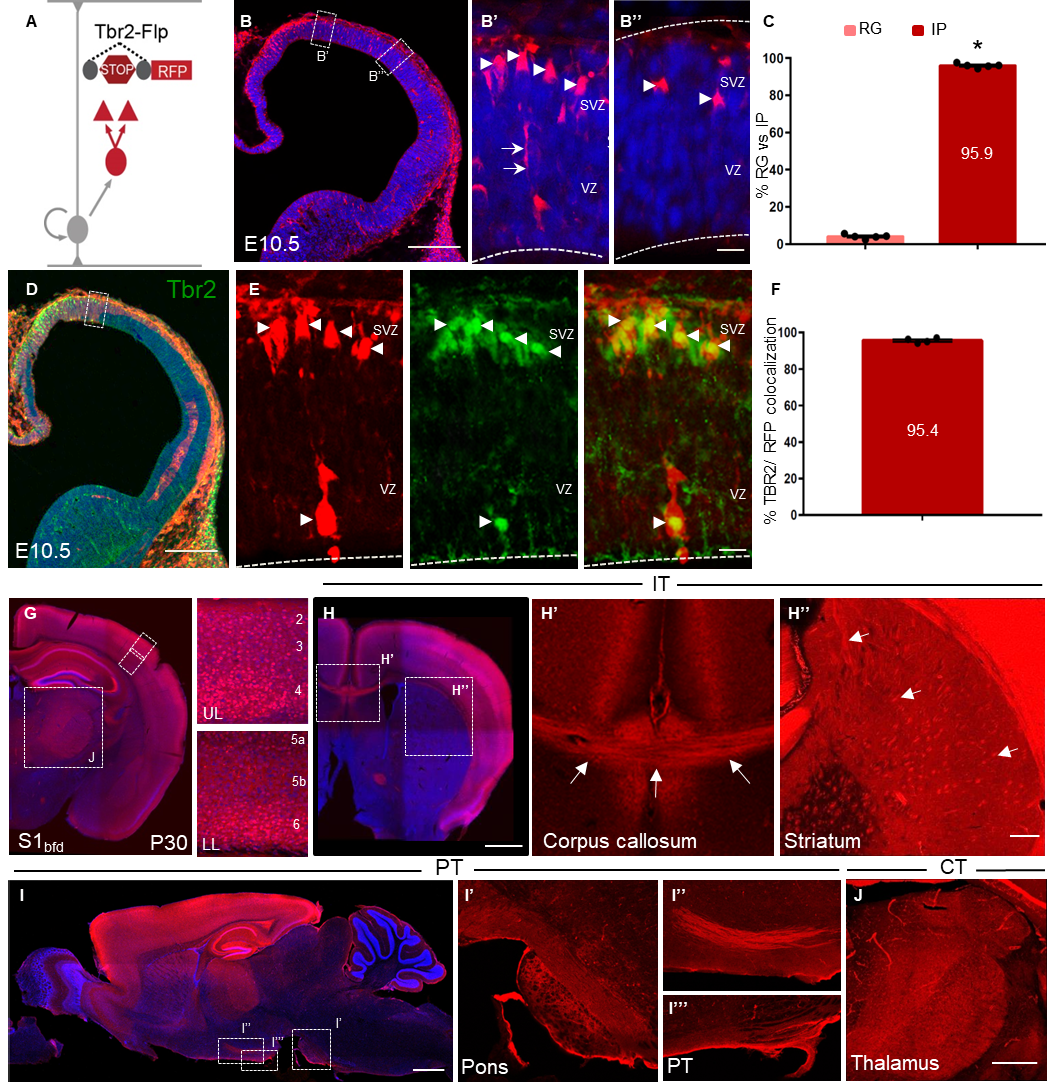

**Supplemental Fig S1: *Tbr2-2A-Flp* captures IPs producing all major cortical PyN classes**

(A) Genetic strategy using *Tbr2-2A-Flp* and a *Flp*-dependent reporter to label all IPs and their progeny. (B) E10.5 *Tbr2* coronal hemisection labels IPs (arrowheads, B’, B’’) in the pallial neuroepithelium. (C) Quantification shows IPs constitute 95.9% of progenitors, with sparsely labeled RG-like cells (arrows, B’). (D) Representative coronal hemisection from E10.5 *Tbr2* brain using the strategy in (A,B) with anti-Tbr2 immunohistochemisty (green). (E) High magnification views from inset in (D) showing tdtomato (red, left), anti-Tbr2 (green, middle) and merge (right). Arrowheads indicate double-labeled cells. The occasional labeling of apical progenitors at this stage are also TBR2 positive. The absence of prominent radial fibers in these apical progenitors suggest that they are unlikely RGs and more likely short neural precursors (SNPs) [S1]. (F) Quantification shows 95.4% tdTomato cells are labeled with Tbr2. (G) P30 coronal hemisection labels PNs across cortical laminae in upper (UL) and lower layers (LL) (high magnification, right). (H) Anterior coronal section reveals that iPNs project across the corpus callosum (H’; arrows) and to the striatum (H’’; arrows). (I) Sagittal section shows cellular RFP expression restricted largely to cortical structures. iPNs project to the pons (I’) and along the pyramidal tract (PT, I’’, I’’’). (J) High magnification view of (G) reveals iPN axons labeled in the thalamus. Quantification in (C) was done from 70-100 cells in 5 embryos each, 2 litters. Mean values are number of cells ± SEM. *P<0.0001 (compared to RG), unpaired Student’s t-test. Quantification in (F) was done from 100 cells in 4 embryos each, 2 litters. Ventricle and marginal zone indicated by dashed lines (B’,B’’,E). Scale bars, low mag 100μm (B,E); high mag 20μm (B’,B’’,D), 1mm (G,H,I), 200μm (J). Abbreviations: RG, radial glial cell; IP, intermediate progenitor; VZ, ventricular zone; SVZ, subventricular zone; IT, intratencephalic, PT, pyramidal tract; CT, corticothalamic; S1_bfd_, primary somatosensory barrel field cortex; UL, upper layer; LL, lower layer. Related to Figure 1.

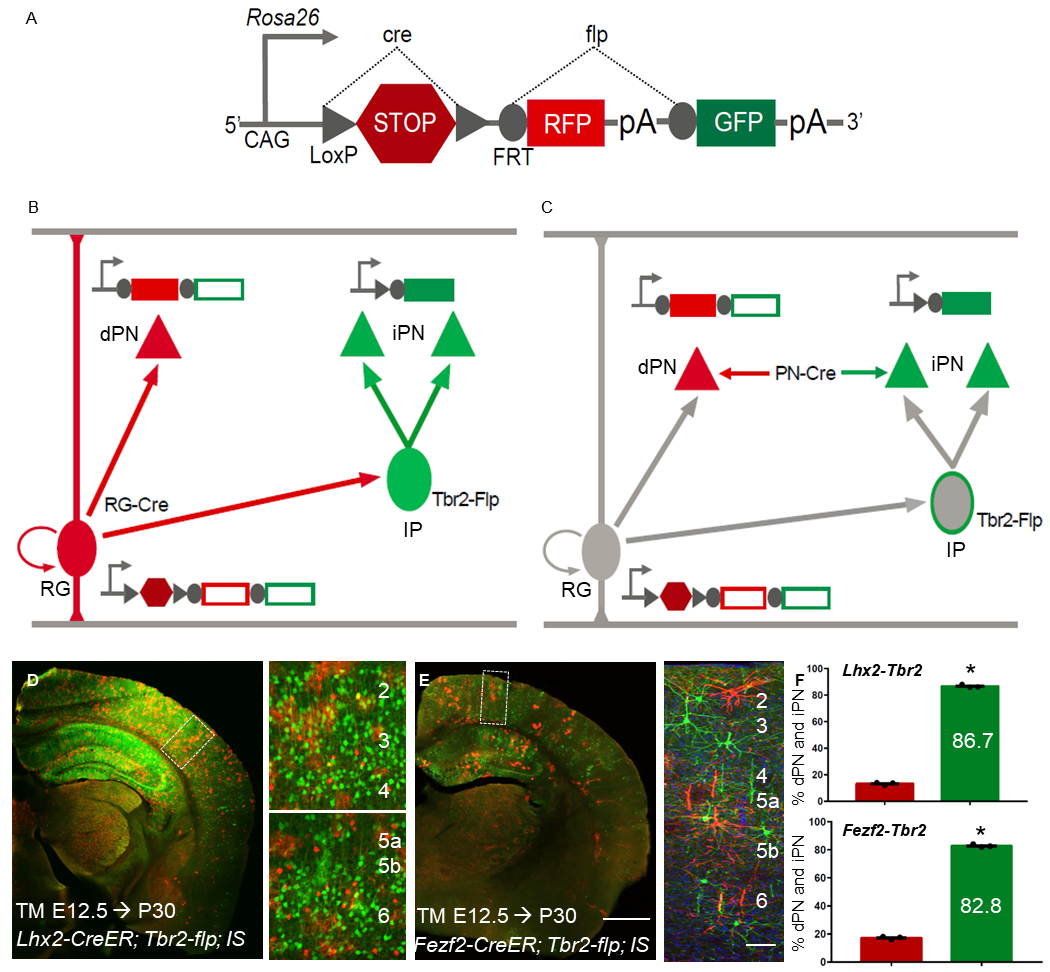

**Supplemental Fig S2: The use of intersection-subtraction strategy for fate mapping dNG and iNG.**

(A) Schematic of the Intersection-Subtraction reporter allele. The transcription and translation STOP cassette contains a SV40 polyA sequence. Cre recombination removes the STOP and activates RFP expression. Cre and Flp recombination further removes the RFP cassette and activates GFP expression. GFP expression can be activated by either Cre-then-Flp or Flp-then-Cre sequential recombinations. (B,C) Within the cortical neuroepithelium, dNG mediated by RGs (red) and iNG through IPs (green) generate PNs (triangles) revealed by the IS strategy (A).

(B) Fate mapping from RGs using driver lines expressing Cre in RGs. Cre recombination in a RG generates an activated RFP expression cassette, which is then transmitted to its progeny, either a PN or an IP. A subsequent Flp recombination in IPs expressing *Tbr2-Flp* then removes the RFP cassette and generates an activated GFP cassette, which is transmitted to the two IP-derived PNs.

(C) Fate mapping using driver lines expressing Cre in postmitotic PNs. Along the dNG pathway, Cre expression in postmitotic PNs removes STOP and activates RFP expression in dPNs. Along the iNG pathway, constitutive *Tbr2-Flp* expression in IPs first removes the RFP cassette in all IP-derived iPNs at the embryonic stage; subsequent postmitotic Cre combination in a subset of iPNs defined by PN-Cre will remove the STOP and activate GFP expression.

(D-F) Different *RG-CreER* driver lines, when combined with IS reporter line and *Tbr2-Flp,* can simultaneously resolve dNG and iNG derived subpopulations. (D) *Lhx2^+^* RGs generate dPN (RFP) and iPN (GFP) across all cortical layers in *Lhx2-CreER; Tbr2-Flp; IS* mice induced at E12.5. High magnification views (right) show the vast majority of PNs are produced through iNG. (E) *Fezf2^+^*RGs also generate dPN (RFP) and iPN (GFP) across all cortical layers in *Fezf2-CreER; Tbr2-Flp; IS* mice induced at E12.5. High magnification views (right). (F) Quantification shows 86.7% Lhx2-derived (top) and 82.8% Fezf2-derived (bottom) PNs are generated from iNG. 1000 cells were counted in 3 mice for (D, F top) and 300 cells were counted in 3 mice for (E, F bottom), mean values are number of cells ± SEM. *P<0.0001 (compared to RFP), unpaired Student’s t-test. Scale bars (D,E), low mag 1mm, high mag 200μm. Abbreviations: RG, radial glial cell; IP, intermediate progenitor; PN, projection neuron. Related to Figures 1, 3.

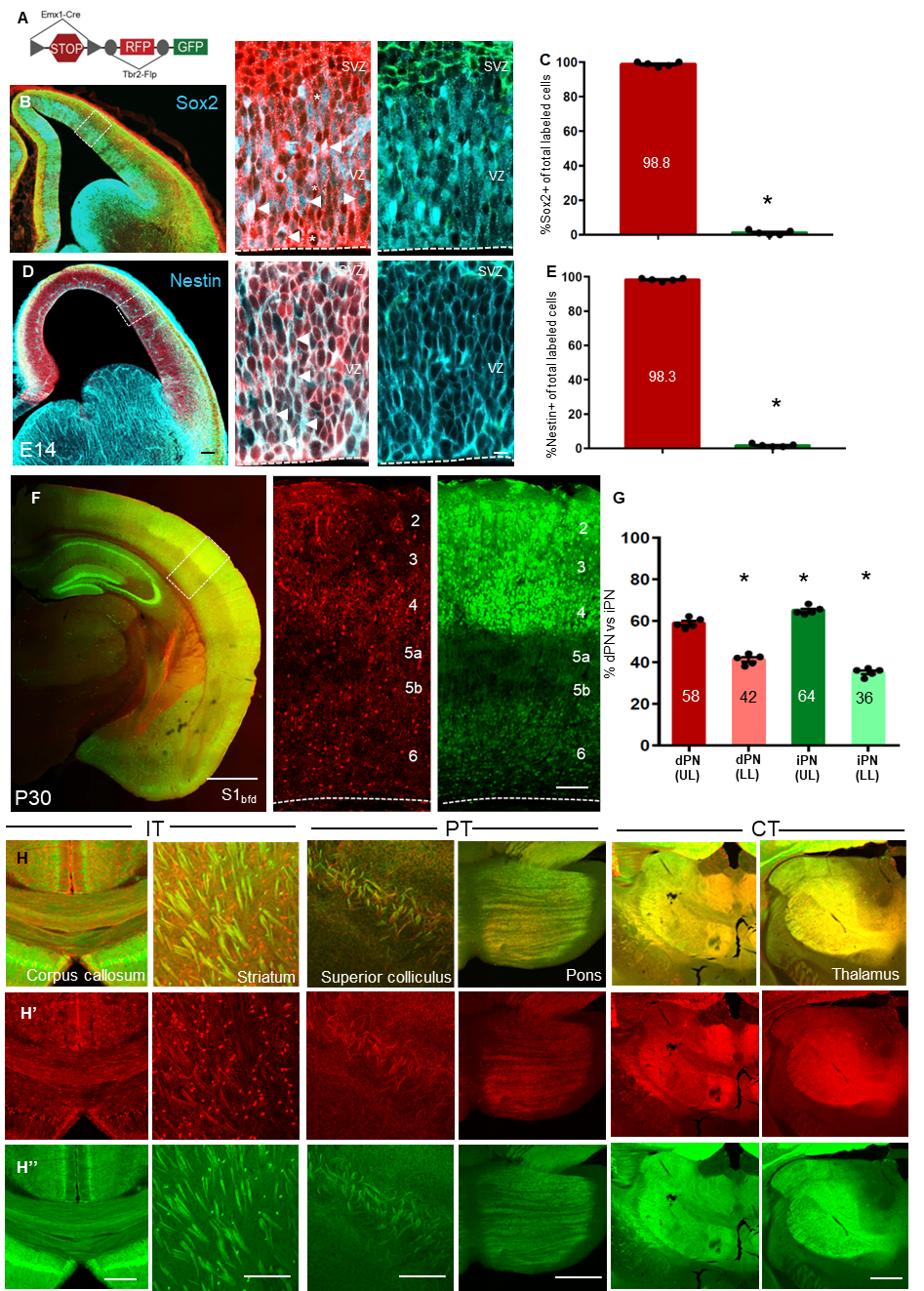

**Supplemental Fig S3: dNG and iNG generate all major classes of PNs in the neocortex**

(A)  IS strategy in combination with *Emx1-Cre* and *Tbr2-Flp* to label RGs and IPs in the cortical neuroepithelium and dPNs (RFP) and iPNs (GFP) in the neocortex (also see Fig.1). (B) Coronal hemi-sections from E14 cortical neuroepithelium with strategy described in (A), labeled with antibodies for RG marker, Sox2. High magnification images (right) show colocalization in the VZ (arrowheads). Some RGs (RFP) are not labeled with Sox2 (asterisks). (C) Quantification shows 98.8% of labeled cells (RFP/GFP) that colocalize with Sox2 are RGs (RFP). (D) Coronal hemisections similar to (B) labeled with anti-Nestin antibodies, another RG marker shows colocalization with RFP(+) RGs (arrowheads) in high magnification views (right). (E) 98.3% of total labeled cells that are Nestin^+^ are RGs (RFP). (F) Fate mapped PNs^Emx1^ from dNG and iNG are present across all cortical laminae (high magnification, right). (G) Quantification showing the distribution of dPNs in UL (58%) and LL (48%) as well as iPNs in UL (64%) and LL (36%) in S1_bfd_ of the neocortex. (H) Both dPNs (H’) and iPNs (H’’) in the neocortex project to all major cortical and subcortical targets: across the corpus callosum, striatum (IT); superior colliculus, pons (PT) and the thalamic nuclei (CT). 200 cells were counted in 5 embryos each from 2 litters, data are mean ± SEM. *P<0.0001 (compared to GFP), unpaired Student’s t-test (C,E).1000 cells were counted in 5 mice, data are mean ± SEM. *P<0.0001 (compared to dPN, UL) one way ANOVA (G). Scale bars, 100μm low mag, 20μm high mag (B,D), 1mm (F), 200μm (all other scale bars). Abbreviations: UL, upper layers (layers 2-4); LL, lower layers (layers 5-6); S1_bfd_, primary somatosensory barrel field cortex; RG, radial glial cell; IP, intermediate progenitor; VZ, ventricular zone; SVZ, subventricular zone. Related to Figure 1.

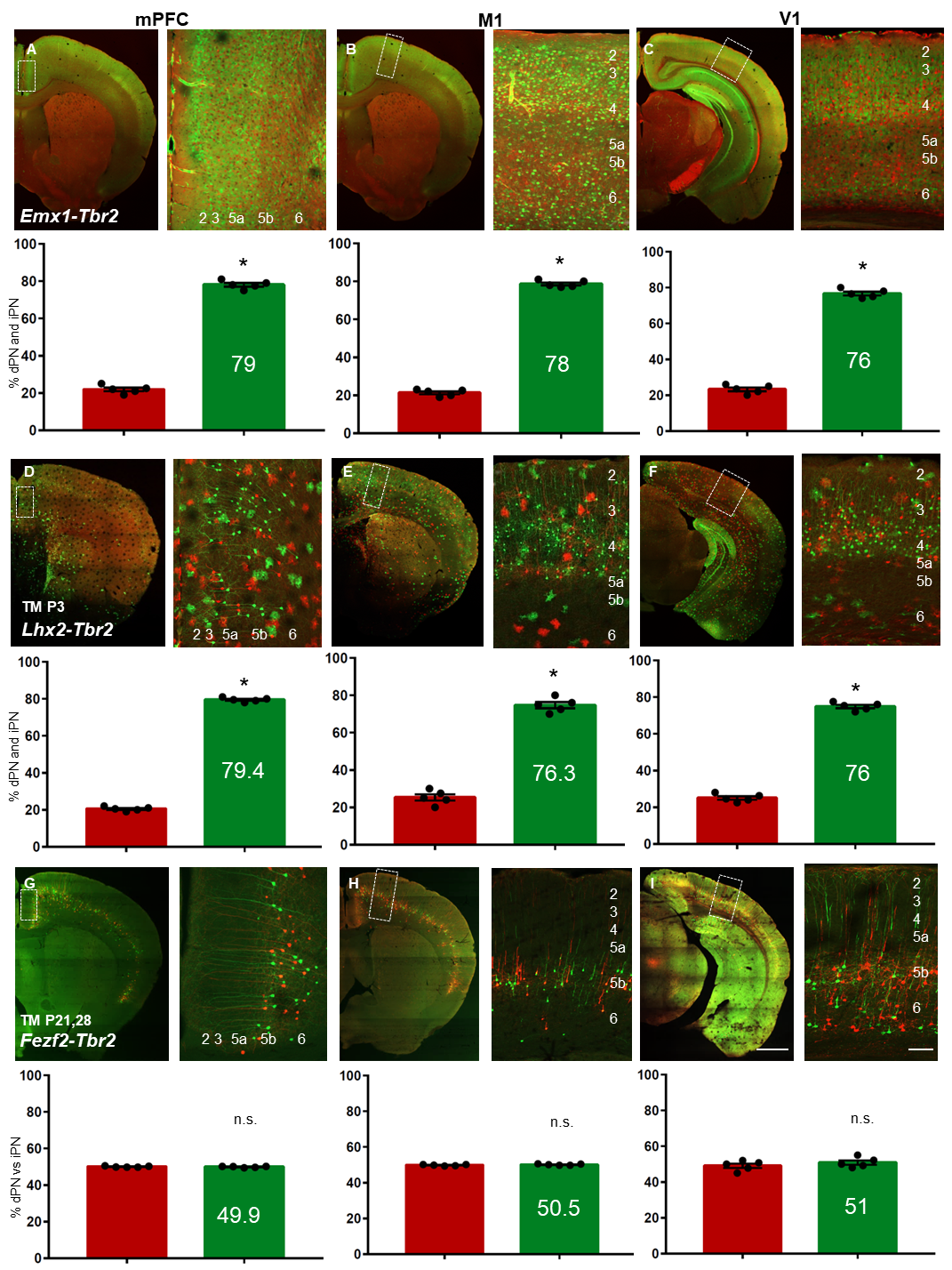

**Supplemental Fig S4: dNG and iNG differentially contribute to neocortical PN projection types across different cortical areas**

Representative coronal hemisections and quantifications of dPN and iPN from the neocortex of P30 *Emx1-Cre* (A-C), P3 tamoxifen induced *Lhx2-CreER* (D-F) and P21 tamoxifen induced *Fezf2-CreER* (G-I) driver lines combined with *Tbr2-Flp* and IS reporter across different cortical areas; mPFC (A,D,G), M1 (B,E,H) and V1 (C,F,I). Percentage iPNs generated are indicated on the bar graphs. 750-1500 cells were counted in 5 mice, data are mean ± SEM. *P<0.0001 (compared to dPN) (A-F), P(n.s.) = 0.598 (G), P(n.s.) = 0.2841 (H), P(n.s.) = 0.3439 (I) unpaired Student’s t-test. Scale bars 1mm (low mag), 100μm (high mag). Abbreviations: mPFC, medial prefrontal cortex; M1, primary motor cortex; V1, primary visual cortex; dPN, directly born projection neurons; iPN, indirectly born projection neurons. Related to Figures 1, 3.

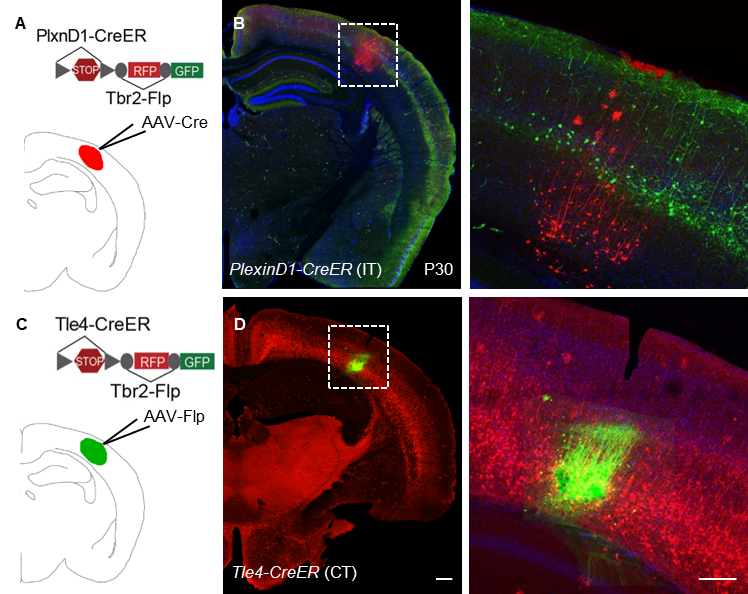

**Supplemental Fig S5: Control for IS reporter functionality to confirm that PNs^Plexind1^ are iNG derived and PNs^Tle4^ are dNG derived**

(A) Schematic of *AAV-Cre* injection in S1_bfd_ of an adult *PlxnD1-CreER;Tbr2-lp;IS* mice.

(B) Whereas *PlxnD1-CreER* activated GFP expression in IP ^Tbr2^-derived PNs, a generic CMV promoter-driven AAV-Cre activated RFP expression in S1_bfd_, indicating that the RFP cassette is intact in the IS reporter.

(C) Schematic of *AAV-Flp* injection in layer 6 in S1_bfd_ of *Tle4-CreER;Tbr2-Flp;IS* mice.

(D) Whereas *Tle4-CreER* activated RFP expression, a generic CMV promoter-driven AAV-Flp activated RFP expression in layer 6 of S1_bfd_, indicating that the GFP cassette is intact in the IS reporter. Scale bars 1mm (low mag); 100μm (high mag). IT, intratencephalic, CT, corticothalamic. Related to Figure 3.

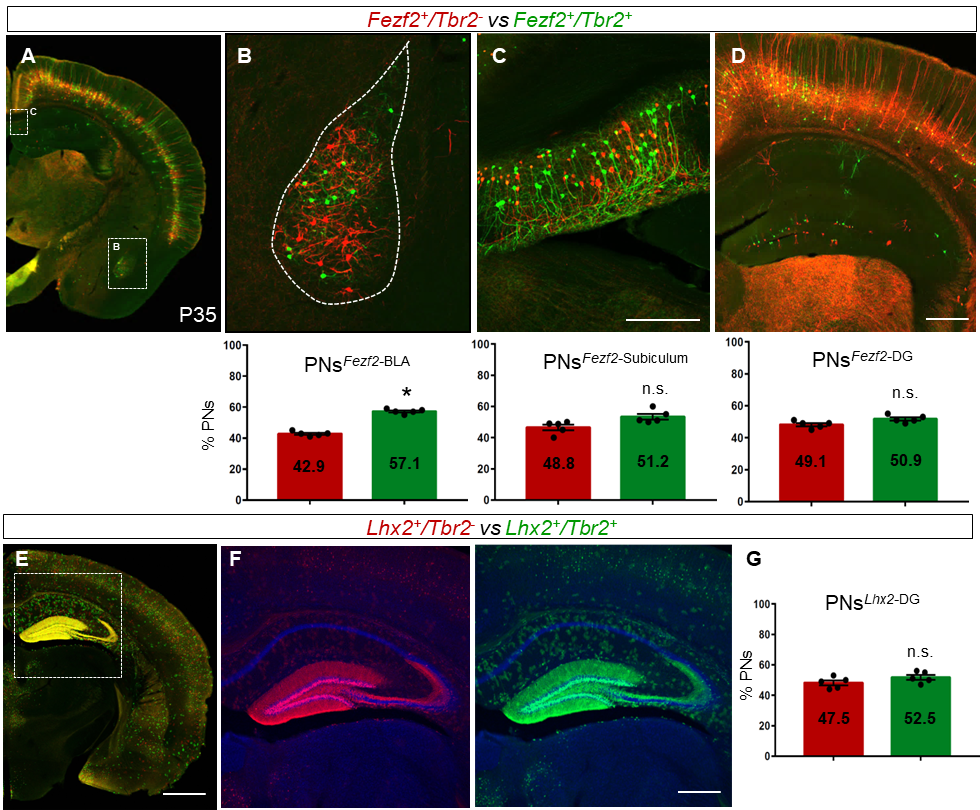

**Supplemental Fig. S6: dNG and iNG contribute differentially to PNs^Fezf2^ and PNs^Lhx2^ in cortical structures**

(A) P21-induced *Fezf2-CreER; Tbr2-flp; IS* coronal hemisection shows differential labeling of dNG and iNG derived PNs^Fezf2^ in different cortical structures at P35. (B) In the BLA, 42.9% PNs^Fezf2^ are dNG-derived and 57.1% are iNG-derived. (C) 48.8% subiculum PNs^Fezf2^ are born via dNG and 51.2% are iNG-generated. (D) The sparsely labeled DG shows a distribution of 49.1% dNG-derived and 50.9% iNG-derived PNs^Fezf2^. (E) *Lhx2-CreER;Tbr2-Flp;IS* induced at P3 labels the DG. (F) PNs*^Lhx2^* are produced via both dNG (RFP) and iNG (GFP), as seen from the higher magnification DG images. (G) Quantification reveals dNG and iNG in the DG are distributed 47.5% and 52.5% respectively. For quantifications, 6-10 sections each from 5 animals were counted. Data are mean ± SEM. *P<0.0001 (compared to dPN, B), *P<0.03 (compared to dPN, C), *P<0.0394 (compared to dPN, D), P(n.s.) = 0.1626 (compared to dPN, G), unpaired Student’s t-test. Scale bars, 1mm (A,E); 200μm (B,C,D,F). Abbreviations: BLA, basolateral amygdala; DG, dentate gyrus. Related to Figures 3,4.

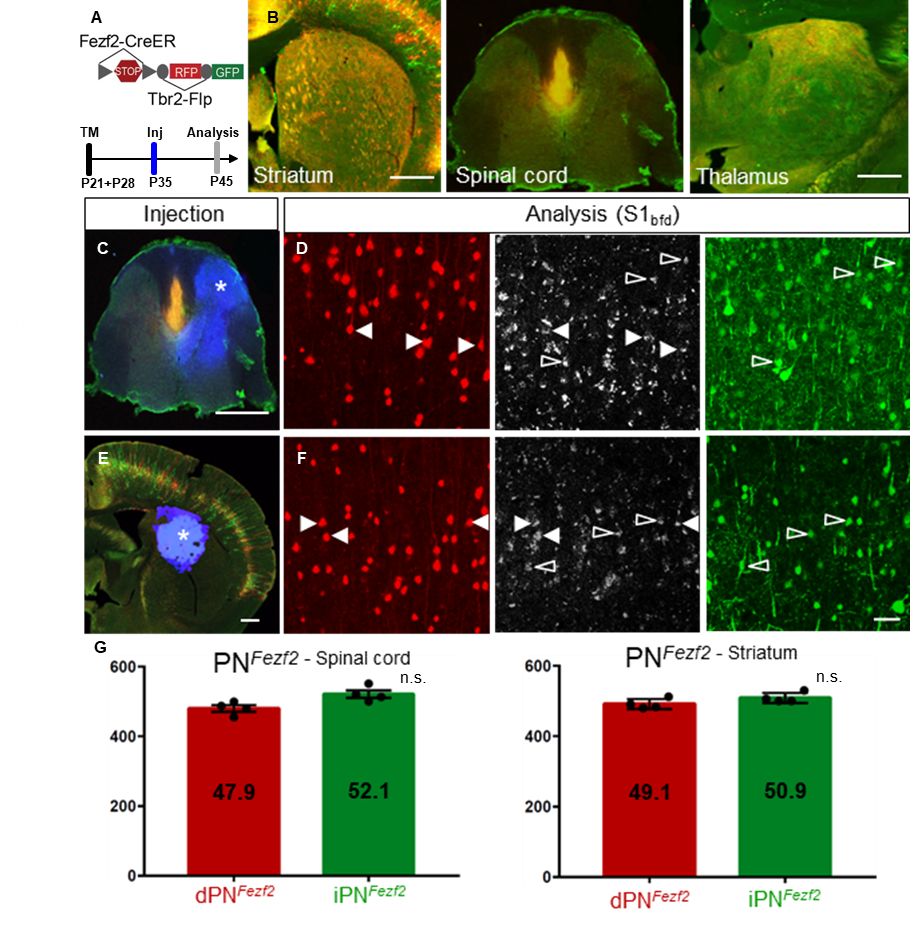

**Supplemental Fig. S7: dPNs^Fezf2^ and iPNs^Fezf2^ project to the spinal cord and striatum in near equal proportions**

(A) (Top) *Fezf2-CreER;Tbr2-Flp;IS* strategy to label dPNs (RFP) and iPNs (GFP). (Bottom) Experimental scheme shows TM induction at P21 to label PNs^Fezf2^, followed by CTB^647^ injection at P35 to retrogradely label projection targets. Brains were analyzed at P45. (B) dPNs^Fezf2^ and iPNs^Fezf2^ both project to the striatum, the spinal cord and the thalamus. (C) CTB was injected in the cervical spinal cord, C1-C4 segments (asterisk).  (D) Analysis in S1_bfd_ to quantify CTB colocalization with dPNs^Fezf2^ (arrowheads, RFP) and iPNs^Fezf2^ (open arrowheads, GFP). (E) CTB injection in the striatum (asterisk). (F) Similar co-localization analysis as in (D). (G) (Left) Quantification spinal cord CTB injection shows colocalization with 47.9% dPNs^Fezf2^ and 52.1% iPNs^Fezf2^. (Right) CTB injection in the striatum shows colocalization with 49.1% dPT*^Fezf2^* and 50.9% iPT*^Fezf2^*. For quantification, ~1000 cells were counted from 5 animals each. Data are mean ± SEM. *P<0.0289 (compared to dPN, G left), P(n.s.) = 0.1355 (compared to dPN, G right), unpaired Student’s t-test. Scale bars, 1mm (C,E); 200μm (B); 100μm (D,F). Abbreviations: S1_bfd_, primary somatosensory barrel field cortex; Inj, injection; TM, tamoxifen induction; Related to Figure 4.

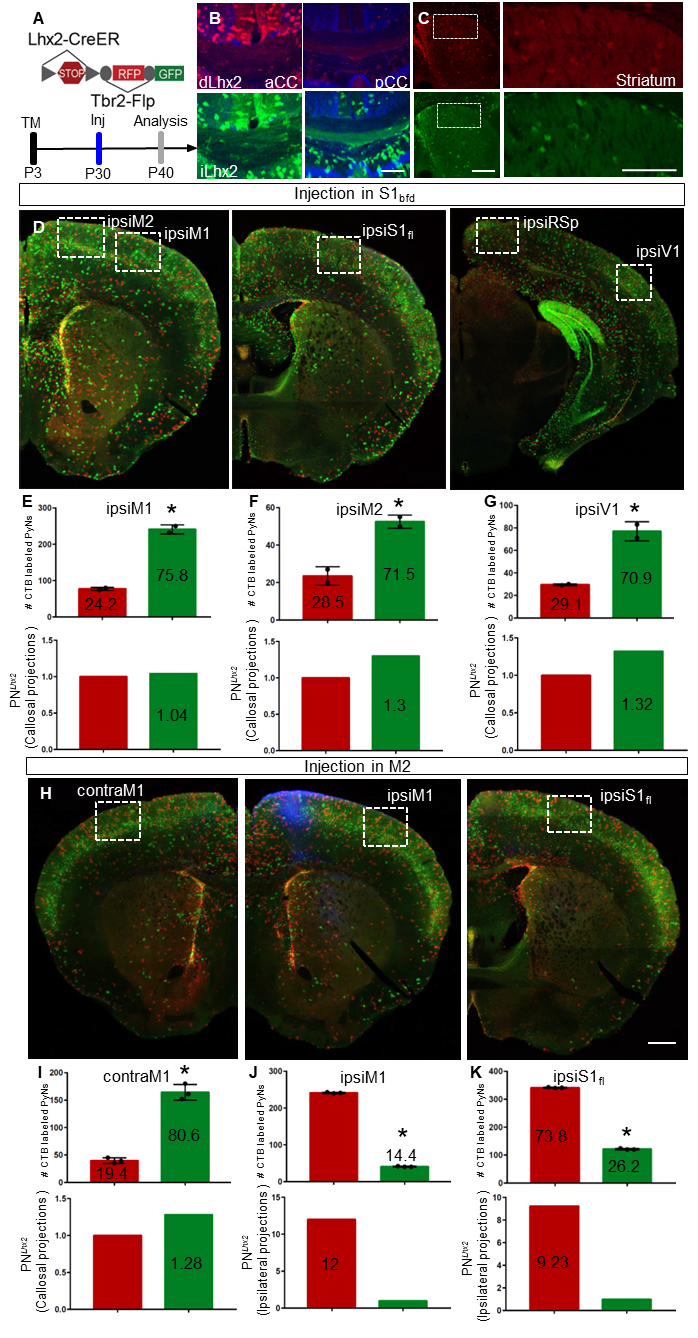

**Supplemental Fig. S8: dPNs^Lhx2^ project preferentially to ipsilateral cortical areas**

(A) (Top) *Lhx2-CreER; Tbr2-Flp; IS* strategy to label dPNs (RFP) and iPNs (GFP). (Bottom) Experimental paradigm with TM induction at P3 to label PNs^Lhx2^, followed by CTB injection at P30. The brains were analyzed at P40. (B) The corpus callosum shows projections from both dPNs^Lhx2^ and iPNs^Lhx2^ at anterior (aCC) and posterior (pCC) levels. (C) PNs^Lhx2^ project sparsely to the striatum. (D) Several cortical areas were analyzed for heterotypic contralateral projections when injected in contraS1_bfd_ from PNs^Lhx2^: ipsiM1, ipsiM2, ipsiS1_fl_, ipsiRsp and ipsiV1 (E) Quantification in ipsiM1 (top) shows that CTB colocalizes with 24.2% dPNs^Lhx2^ and 75.8% iPNs^Lhx2^ (~400 cells). (Bottom) Normalization to the total PNs^Lhx2^ labeled reveals similar projections (1.04-fold difference) from dPNs and iPNs. (F) (Top) Analysis in ipsiM2 reveals CTB colocalization with 28.5% dPNs^Lhx2^ and 71.5% iPNs^Lhx2^ (~90 cells). (Bottom) Normalizing values to the number of PNs^Lhx2^ shows similar projections (1.3-fold difference) between dPNs and iPNs. (G) (Top) CTB colocalizes with 29.1% dPNs^Lhx2^ and 70.9% iPNs^Lhx2^ in V1 (~120 cells). (Bottom) There are similar projections (1.32-fold difference) from dPNs and iPNs upon normalization. No colocalization seen in ipsiS1_fl_ and ipsiRSp (H) ipsiM1, contraM1 and ipsiS1_fl_ were analyzed for projections to M2. (I) Analysis in contraM1 shows colabeling with 19.4% dPNs^Lhx2^ and 80.6% iPNs^Lhx2^ (top, ~200 cells) with no difference (1.28-fold) between dPNs^Lhx2^ and iPNs^Lhx2^ (bottom). (J) ipsiM1 (top) shows that CTB colabels with 85.6% dPNs^Lhx2^ and 14.4% iPNs^Lhx2^ (~300 cells) showing 12-fold higher projections from dPNs^Lhx2^ (bottom). (K) Quantification in ipsiS1_fl_ shows colocalization with 73.8% dPNs^Lhx2^ and 26.2% iPNs^Lhx2^ (top, ~500 cells) resulting in 9.23-fold higher projections from dPNs^Lhx2^ (bottom).

Quantification was done from 3-4 animals each. Data are mean ± SEM. *P<0.0001 (compared to dPN) unpaired Student’s t-test. Scale bars, 200um (A,B); 1mm (D,H). Abbreviations: Inj, injection; TM, tamoxifen induction; aCC, anterior corpus callosum; pCC, posterior corpus callosum; ipsiM1, ipsilateral primary motor cortex; ipsiM2; ipsilateral secondary motor cortex; ipsiS1_Fl_, ipsilateral primary somatosensory forelimb cortex; ipsiRSp, ipsilateral retrosplenial cortex; ipsiV1, ipsilateral primary visual cortex; contraM1, contralateral primary motor cortex. Related to Figure 4.
